# Remote-refocusing light-sheet fluorescence microscopy enables 3D imaging of electromechanical coupling of hiPSC-derived and adult cardiomyocytes in co-culture

**DOI:** 10.1101/2023.01.28.526043

**Authors:** L Dvinskikh, H Sparks, L Brito, K MacLeod, SE Harding, C Dunsby

## Abstract

Improving cardiac function through stem-cell regenerative therapy requires functional and structural integration of the transplanted cells with the host tissue. Visualizing the electromechanical interaction between native and graft cells necessitates 3D imaging with high spatio-temporal resolution and low phototoxicity. A custom light-sheet fluorescence microscope was used for volumetric imaging of calcium dynamics in co-cultures of adult rat left ventricle cardiomyocytes and human induced pluripotent stem cell-derived cardiomyocytes. Aberration-free remote refocus of the detection plane synchronously to the scanning of the light sheet along the detection axis enabled fast dual-channel 3D imaging at subcellular resolution without mechanical sample disturbance at up to 8 Hz over a ∼300 µm × 40 µm × 50 µm volume. The two cell types were found to undergo electrically stimulated and spontaneous synchronized calcium transients and contraction. Electromechanical coupling was found to improve with co-culture duration, with 50% of adult-CM coupled after 24 hours of co-culture, compared to 19% after 4 hours (p = 0.0305). Immobilization with para-nitroblebbistatin did not prevent calcium transient synchronization, with 35% and 36% adult-CM coupled in control and treated samples respectively (p = 0.91), indicating that electrical coupling can be maintained independently of mechanotransduction.

## Introduction

Cardiac muscle is thought to have limited natural regenerative capacity, with less than 1% cardiomyocyte turnover rate in adults [1], and is unable to compensate for damage after myocardial infarction (MI) [2], [3]. While stem-cell therapy has shown some promise for regenerative treatment of cardiovascular diseases [4], the marginal improvement in heart function has been attributed to repair-promoting paracrine effects and has been limited by early-stage cell washout and death [5], and an absence of electrical coupling between host and graft cells [6]. Successful electromechanical coupling has previously been demonstrated in non-primate models in the form of synchronized calcium transients and the formation of electromechanical junctions between native heart tissue and grafted cardiomyocytes derived from the pluripotent embryonic stem cells (ESCs) [7]. Induced pluripotent stem cells (iPSCs) are directly derived from somatic cells in adult tissue [8] and can be obtained autologously, reducing the risk of the transplanted cells being rejected by the immune system [9].

Direct support of cardiac contractile function is thought to require structural and functional integration of the graft cells with the host tissue, the lack of which can result in independent contraction of the transplanted cells. During the establishment of coupling, the partial connections between immature iPSC-derived cardiomyocytes (iPSC-CM) and host tissue can cause severe arrhythmia-based complications [10]. The immaturity of induced pluripotent stem cell-derived cardiomyocytes (iPSC-CM) compared to adult cardiomyocytes (adult-CM) is manifested in their differences in structure and calcium handling. A key attribute of iPSC-CM is their automaticity: successfully differentiated cells undergo spontaneous beating that includes a global calcium transient [11]. The limited functionality of iPSC-CM is attributed to their structural immaturity [12]. Compared to adult-CM, iPSC-derived cardiomyocytes lack t-tubule structure and have fewer and inhomogeneously clustered ryanodine receptors (RyR) [13], reducing the efficiency of calcium-induced calcium release (CIRC), resulting in weaker and slower transients [14]-[16].

Observing and characterizing the electromechanical interaction between the graft and host cells is critical to understanding and assessing of the integration of the transplanted cells and the overall potential of stem cell regenerative therapy. At the single-cell level, synchronous contraction of individual cardiomyocytes requires efficient transmission of mechanical forces and electrical signals. In addition to electrical coupling through gap junction formation, mechanotransduction has been shown to regulate the transmission of contractile force between stem cell-derived cardiomyocytes of different origin and different calcium handling and contractile properties [17]. Calcium cycling is a key regulator of excitation-contraction coupling, and hence optical mapping of calcium dynamics in cardiac cells is essential to characterizing the electromechanical coupling between graft and host cardiomyocytes. Imaging calcium dynamics in multicellular culture requires rapid volumetric imaging, ideally with high and isotropic spatial resolution. Light-sheet fluorescence microscopy (LSFM), also referred to as selective plane illumination microscopy (SPIM) [18], enables optically sectioned imaging with low phototoxicity and fast parallelized widefield acquisition, making it a suitable imaging modality for this application.

In this paper, we investigate the dynamics of early coupling between stem cell-derived and adult cardiomyocytes. A dual-objective light-sheet fluorescence microscope is used for 3D imaging of calcium dynamics in a co-culture of spontaneously beating human induced pluripotent stem cell-derived cardiomyocytes (hiPSC-CM) and the usually quiescent adult left ventricle cardiomyocytes isolated from male rats. The novel LSFM system, described in detail in our previous work [19], implements folded aberration-free remote refocus [20], [21] of the detection plane, enabling fast volumetric imaging without any direct mechanical sample disturbance. We have previously used the remote-refocusing LSFM system for proof-of-concept 3D live cell imaging in adult ventricular cardiomyocytes [19]. In this paper we implement the novel LSFM system for dual spectral channel imaging at up to 8 Hz over a ∼300 µm × 40 µm × 40 µm field of view (FOV) at subcellular resolution. We demonstrate the potential of this system to image the coupling dynamics in heterogenous cardiomyocyte co-culture, investigating the influence of co-culture duration and motion uncoupling on the co-culture efficiency.

## Methods

### Sample preparation

All studies were carried out with the approval of the local Imperial College London ethical review board and the Home Office, UK and in accordance with the Animals (Scientific Procedures) Act 1986 Amendment Regulations 2012, and EU directive 2010/63/EU, which conforms to the Guide for the Care and Use of Laboratory Animals published by the U.S. National Institutes of Health under assurance number A5634-01. The reporting in the manuscript follows the recommendations in the ARRIVE (Animal Research: Reporting of *In Vivo* Experiments) guidelines.

#### hiPSC-CM differentiation and culture

The hiPSC-CM cells were differentiated from the human lung fibroblast cell line IMR-90 (iPS(IMR90)-4, WiCell) following the protocol outlined in [22]. From day 15 after the start of differentiation, the hiPSC-CM were maintained in *hiPSC-CM maintenance media* containing RPMI 1640 (Thermo Fisher Scientific) with added 1× B27 (Thermo Fisher Scientific) and 1% AA (Antibiotic-Antimycotic, Thermo Fisher Scientific). The replating was conducted on day 23 after the start of differentiation using the following process. First, cell detachment was attained using a cell dissociation solution (at 2:2:1 ratio, Cell Dissociation Buffer (Thermo Fisher Scientific): RPMI 1640: 0.05% Trypsin-EDTA (Thermo Fisher Scientific)) for 15 min at 37°C. Next, trypsin de-activation was achieved using the hiPSC-CM maintenance media supplemented with 10% FBS (Fetal Bovine Serum, Thermo Fisher Scientific), and cells were pelleted by centrifugation at 200 g for 5 minutes. The cells were replated at 150 k/cm^2^ cell density onto plastic coverslips (Nunc Thermanox 150067, Thermo Fisher Scientific) in hiPSC-CM maintenance media with added 10% FBS and 10 μM Rock inhibitor Y-27632 (Stratech). Prior to replating, the plastic coverslips had been cut to fit within the 1 cm^2^ area chambers in a µ-Slide 8-Well Glass-Bottom Chamber (ibidi), and coated with fibronectin from bovine plasma (Sigma-Aldrich), diluted in 1:100 in 1× PBS for 1 h at 37°C. One day after replating, the media was replaced by the hiPSC-CM maintenance media and maintained at 5% CO_2_, 37°C, until at least 4 weeks past start of differentiation prior to use in experiments, with media change every 3 days.

#### hiPSC-CM co-culture with adult-CM

The hiPSC-CM were co-cultured with left-ventricle cardiomyocytes, freshly isolated following the method outlined in [23] from male rats, anesthetized with 5% isoflurane prior to sacrifice. The cardiomyocytes were suspended in low Ca^2+^ enzyme solution, containing (in mM) NaCl (120), KCl (5), MgSO_4_ (5), Na pyruvate (5), glucose (5), taurine (20), HEPES (10), CaCl_2_ (0.2), pH 7.4 adjusted with 1M NaOH. The adult-CM were re-suspended at 1:10 dilution in M199+ culture medium containing (per 500 ml M199): 5 mL P/S, 1 g Bovine Serum Albumin (BSA, Sigma-Aldrich), 0.33 g creatinine (Sigma-Aldrich), 0.33 g taurine (Sigma-Aldrich), 0.0088 g L-ascorbic acid (Scientific Laboratory Supplies), 0.161 g carnitine hydrochloride (Sigma-Aldrich). To make the co-culture, the RPMI cell media was aspirated almost completely from the hiPSC-CM culture and quickly replaced with the same amount of M199+ culture medium. Next, suspended isolated adult-CM were added dropwise on top of the stem cell culture to achieve a scattered adult-CM distribution (**Supplementary Figure 1a**). The cells were co-cultured at 5% CO_2_, 37°C for varying durations (4 hours, 1 day, 2 days) up until the point of imaging, with daily media changes.

#### hiPSC-CM and co-culture preparation for live fluorescence imaging

The live-cell dual labelling preparation protocol was based on that used by [24] for OPM-based imaging of live cardiomyocytes, and on that used for our proof-of-concept live cell LSFM imaging, described in [19]. For simultaneous monitoring of the calcium dynamics and the cell membrane and microstructure, the cells were dual-labelled with cell-permeant calcium indicator Fluo-4 AM (Thermo Fisher Scientific) and CellMask Orange (CMO) plasma membrane stain (Thermo Fisher Scientific). Using tweezers, the plastic coverslips were carefully transferred into a larger chamber and resuspended in 1 mL M199+ culture medium. The co-cultures were incubated with 0.16% pluronic acid and 5 µM Fluo4-AM in DMSO for 15 minutes at 37°C, followed by the addition of CellMask Orange (CMO) at 1µM for another 5 min. Throughout the incubation period, the chamber containing the coverslip-seeded cells was placed on a rotary mixer (at the lowest speed setting) to ensure homogenous dye distribution and protected from light by wrapping the dish in foil. After a total 20 minutes of incubation, the sample was resuspended in M199+ culture medium, pre-heated to 37°C.

Mechanical contraction was prevented in some samples by decoupling the cell contraction from the calcium dynamics using para-nitroblebbistatin (NBleb, Axol Bioscience), a non-phototoxic low fluorescence myosin inhibitor [25], added to the M199+ and normal Tyrode (NT) solution at 25 µM concentration. After allowing 20 minutes for Fluo-4 AM de-esterification, using tweezers, the coverslip was gently transferred to the imaging chamber, and placed between two stimulating electrodes. The pacing and imaging chamber (**Supplementary Figure 1b**) consisted of a glass slide cut to a rectangle with dimensions of ∼6 × 5 cm, with the electric field stimulation at 1.5× threshold voltage (found to be approximately 20 V), 2 ms pulse duration and 0.5 Hz frequency (unless stated otherwise) delivered using two parallel 4 cm-long electrodes made from 0.8 mm diameter platinum wire (Goodfellow). The electrodes were attached to the glass slide using epoxy adhesive, separated by ∼1 cm and parallel over ∼2 cm. During imaging, the sample, electrodes and front lenses of the two objectives were immersed in 1 mM Ca^2+^ NT solution, pre-heated to 37°C, with liquid immersion maintained by surface tension.

### Data acquisition

The acquisition parameters and imaging modes for LSFM-based acquisition are summarized in **Supplementary Table 1**.

#### Inverted SPIM system with aberration-free remote refocusing

The optical design and operation of the dual-channel SPIM system has been described in detail in [19], and the relevant functionality is briefly outlined here. Fluorescence excitation is achieved using angularly dithered Gaussian light-sheet illumination at a ∼37 degree angle with respect to the horizontal sample surface (**Figure 1a**). A cylindrical lens is used to focus the laser excitation to create a line in the back focal plane of the water immersion 10× 0.3NA illumination objective.

**Figure 1.**
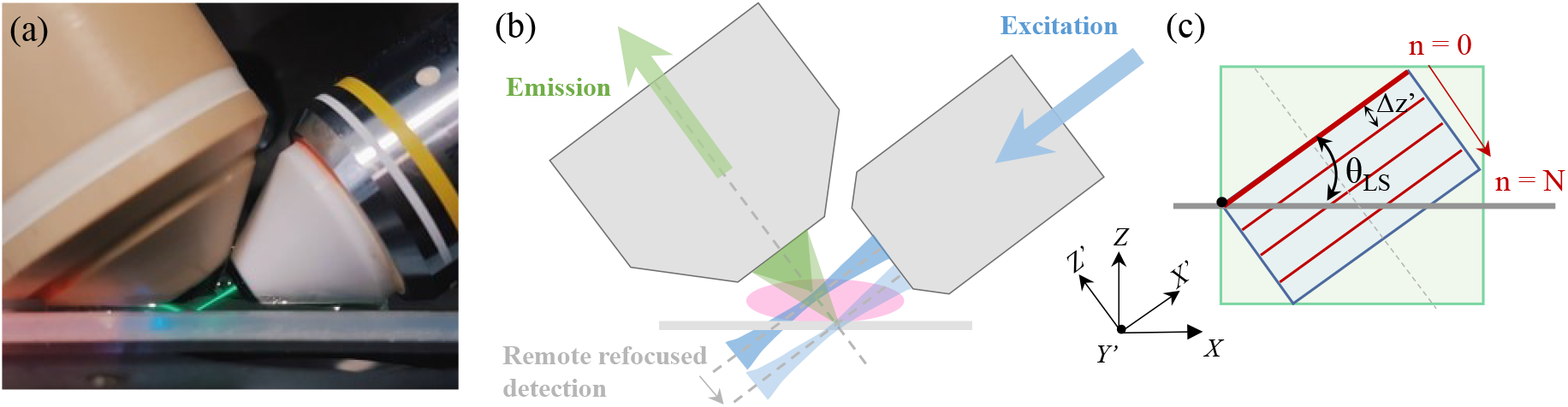
Acquisition geometry for remote-refocusing dual-objective LSFM. (a) Detection (left) and illumination (right) objective geometry, with fluorescein emission used to visualize the light sheet. (b) Schematic of illumination and detection geometry, with the remote refocused detection synchronized with the scanning of the light sheet. (c) Remote refocusing volumetric acquisition geometry, with the focus swept along the Z’ detection axis (dashed grey line), with consecutive images separated by distance Δ*z’* along the detection axis, with the sample remaining stationary. The light-sheet angle with respect to the horizontal surface is indicated by θ_LS_.

Fluorescence emission is collected by a 1 NA 20× water immersion objective, with its optical axis orthogonal to the light-sheet. High-speed volumetric imaging is achieved using folded remote refocusing (**Figure 1b-c**), with the intermediate image formed by the first microscope re-imaged by a second microscope relay consisting of a tube lens and a 0.75 NA 20× air objective onto a remote-refocusing mirror, mounted on a piezoelectric actuator for rapid axial scanning of the detection plane. Unlike stage or detection objective scanning methods, there is no mechanical perturbation to the sample, as the scanning is performed remotely. The system performance compared to other approaches was reviewed in our previous publication [19]. For aberration-free remote refocus, the optical elements are selected such that the lateral and axial magnification from the sample to the remote refocus space is equal to the ratio of their respective refractive indices [20], [21]. Fluorescence is coupled in and out of the folded remote refocusing unit using a quarter waveplate and polarizing beam splitter. For fluorophores with a low steady state anisotropy, the emission is unpolarized and half the light is lost in the polarizing beam splitter, reducing the overall collection efficiency. For fluorescence from randomly oriented fluorophores with a high stead-state anisotropy, the emission is partially polarized and ∼¾ of the fluorescence will pass through the polarizing beam splitter.

The refocused detection plane is reimaged onto a two-dimensional adjustable slit which is used to define the boundary of the FOV, and then reimaged onto a sCMOS detector, with a final sample-to-image magnification of 44×. Simultaneous dual-channel imaging of Fluo-4 and CMO fluorescence was implemented by separating the emission using a 560 nm edge long-pass dichroic beam splitter with 525/50 nm and 630/75 nm emission filters. The image from each spectral channel was directed onto each half of the camera sensor using positioning mirrors. Longitudinal chromatic aberration introduced by our custom tube lens (labelled as “P9” in [19]) in the remote refocusing path resulted in a ∼10 mm axial separation between the image foci of the two spectral channels. This was corrected by introducing two doublets of near-equal but opposite focal length (160 mm and -150 mm) in the Fluo-4 emission path, with principal planes separated by ∼15 mm, which introduced 0.9× magnification of the Fluo-4 spectral channel (See **Supplementary note 1** and **Supplementary Figure 2**)

For 3D imaging, the Gaussian light-sheet was galvo-scanned along the detection axis, synchronously with remote refocusing of the detection plane (see **Supplementary Figure 3** for hardware timing), without any mechanical disturbance of the sample. The actual z-position of the piezo which translates the remote-refocusing mirror was recorded using a capacitive sensor integrated into the actuator. This recorded signal was used as reference for adjusting the driving voltage profile applied to the piezo to achieve the desired scan range, and for determining the linear part of the scan range for acquisition (see **Supplementary Figure 4**). The recorded piezo position was also used during the calibration procedure for determining the GZ galvo voltage required at each piezo actuator position to ensure synchronisation between the axial scanning of the light sheet and the remotely refocused detection plane. The factor limiting the speed for this acquisition mode was the frame rate for a given vertical extent of the camera sensor used to image the selected FOV.

#### 2D LSFM imaging of live hiPSC-CM

For 2D LSFM stem cell imaging, Fluo-4 and CMO were excited simultaneously by the 488 nm and 561 nm laser lines at 25 µW and up to 35 µW power respectively, measured in the back focal plane (BFP) of the excitation objective, with angular dithering of the light sheet within the illumination plane. Fluorescence emission was acquired in the form of two-dimensional dual-channel time-lapse data using a Hamamatsu Orca Fusion sCMOS camera with a 6.5 μm × 6.5 μm pixel size. The camera was operated in normal area external trigger mode, with synchronous readout trigger. Each acquisition had a duration of 57 s, imaged at 175 fps for a 1152 pixel × 512-pixel region of interest (ROI), which corresponds to a 170.2 μm × 38 μm FOV (per spectral channel) in sample space at 44× magnification and 0.1477 μm pixel size. For paced acquisitions, the cells were electrically stimulated starting 1 minute before each acquisition.

#### 3D LSFM imaging of live hiPSC-CM and co-culture

For 3D LSFM imaging, the fluorescence of both CMO and Fluo-4 was excited using only the 488 nm excitation line with an excitation power of 312 µW (measured in the BFP of the excitation objective). For co-culture imaging, sample navigation and pre-find were done in transillumination, with individual adult cardiomyocytes distinguished from the underlying stem cell layer due to their stronger scattering (**Supplementary Figure 5**). Fields of view were selected such that each one contained an adult-CM with the longer axis approximately perpendicular to the light-sheet propagation direction. Each FOV containing an adult cardiomyocyte was imaged first without electrical pacing. Then, after pre-pacing the sample for 1 min prior to the acquisition (2 ms pulse duration, 20 V and 0.5 Hz frequency), imaging was repeated with electrical stimulation. The synchronization monitor signal from the electric field stimulation unit was recorded as a digital input to the DAQ card. The 3D volumes of co-culture data were reconstructed from image stacks consisting of 38 planes with 1.3 µm spacing along the detection axis, with the 15 s duration timelapses acquired at either 120 volumes at 390 fps (8 vps) or 60 volumes at 195 fps (4 vps).

#### Widefield fluorescence and transillumination imaging

For imaging over a larger FOV than achievable with the LSFM system magnification, complementary brightfield transillumination and epi-fluorescence 2D-timelapse of the hiPSC-CM and adult-CM co-culture was carried out on an inverted widefield microscope (IX73, Olympus) equipped with LED illumination (CoolLED pE300) and a B/W CCD digital camera (Hamamatsu C4742-95-12ER). Detection and epi-fluorescence illumination was achieved using infinity corrected air objectives: UplanFL 4×/0.13 NA (1-U2B522, Olympus) and CPlan FL N 10× 0.30 NA PhC (1-U2C543, Olympus). Each acquisition consisted of an image sequence of between 100-200 frames covering durations of 40-60 s, each one with image dimensions of 1344 × 1024 pixels, corresponding to physical image dimensions of 867 × 661 µm and 2167 × 1651 µm for the 10× and 4× objectives respectively. Data was acquired at either 2 fps or 5 fps, and exposure times varied between 5 and 200 ms between the different magnifications and illumination modes. For imaging of calcium dynamics, Fluo-4 was excited using the blue LED (peak at 450nm) and a 472 ± 30 nm (centre and FWHM) excitation filter. Fluorescence was collected through a GFP-3035C-000 filter with a detection range of 520 ± 35 nm. A 495 nm long-pass dichroic mirror was used to separate the excitation and emission.

### Data processing and analysis

#### LSFM data processing, reconstruction and analysis

Co-registration of the two spectral channels for 2D and 3D stem-cell and co-culture data was carried out in MATLAB. First, fixed pattern noise estimated from an average of 1000 frames acquired with the laser shutter closed was subtracted from each raw frame in the dataset. Next, the two spectral channels were split and co-registered using an affine transformation consisting of rotation, two-dimensional scaling, and translation, with Fluo-4 channel images demagnified by a factor of ∼0.9× in each lateral dimension to account for the additional magnification in the Fluo-4 detection path. The exact parameters for the transformation were manually determined by achieving best visible overlap for a dual-channel dataset obtained by imaging 200 nm diameter fluorescent beads (T7280, TetraSpeck™, Thermo Fisher Scientific) embedded in agarose.

For visualization of the 3D-time-lapse data (x-y-z-t), the X’Y image stack, recorded as a sequence of N X’Y frames, where N = Z × T, was re-ordered into a X’Y-z-t image hyperstack in ImageJ. Each X’Y image in the data stack was cropped to the FOV of the light transmitted through the CMO channel to exclude the dark border due to the 2D slit, and axially rescaled to attain a uniform voxel size extending 0.1477 µm in each dimension. Next, the dataset was re-sliced to achieve an X’Z’ view of the sample and rotated by 37° about the y-axis to transform the volume into lab coordinates. The XZ view was resliced to generate XY and YZ views of the sample in lab coordinates. The resliced orthogonal cuts of the volume were rendered as orthogonal slices or maximum intensity projections (MIPs) and the two spectral channels were combined for multicolour visualization. Depth-encoded MIPs along the z-axis of the Fluo-4 channel were generated using the “Z-stack Depth Color Code” Fiji plugin using the “Ice” lookup table [26]. Statistical analysis of significance was carried out in GraphPad Prism 9.1.1 software, with p-values calculated using Chi-square tests, with p < 0.05 considered statistically significant.

## Results

### Imaging calcium dynamics in hiPSC-CM

Live cell imaging of hiPSC-CM culture was performed using 2D- and 3D-timelapse LSFM with and without electric field stimulation. Using high-speed 2D LSFM timelapse imaging, spontaneous calcium transients, and electrically stimulated transients at varying pacing periods (T = 2 s, 4 s and 6 s) were observed in hiPSC-CM culture (**Supplementary Note 2**).

Fluo-4 (a), CMO (b) and merged channel (c) orthogonal cuts through a hiPSC-CM culture imaged in 3D are shown in **Figure 2**, with the corresponding timelapse presented in **Supplementary Video 1**. Formatting…The same FOV was imaged in two consecutive acquisitions, first unpaced, then with electrical pacing at 0.5 Hz. The unpaced acquisition demonstrated spontaneous transients beginning 6.25 s apart, while the paced hiPSC-CM culture had stimulated transients at the paced frequency but at a lower amplitude. This reduced amplitude with increased transient frequency demonstrated in **Supplementary Figure 6** and **Figure 2** is in agreement with the negative force-frequency response characteristic of hiPSC-CM [27].

**Figure 2.**
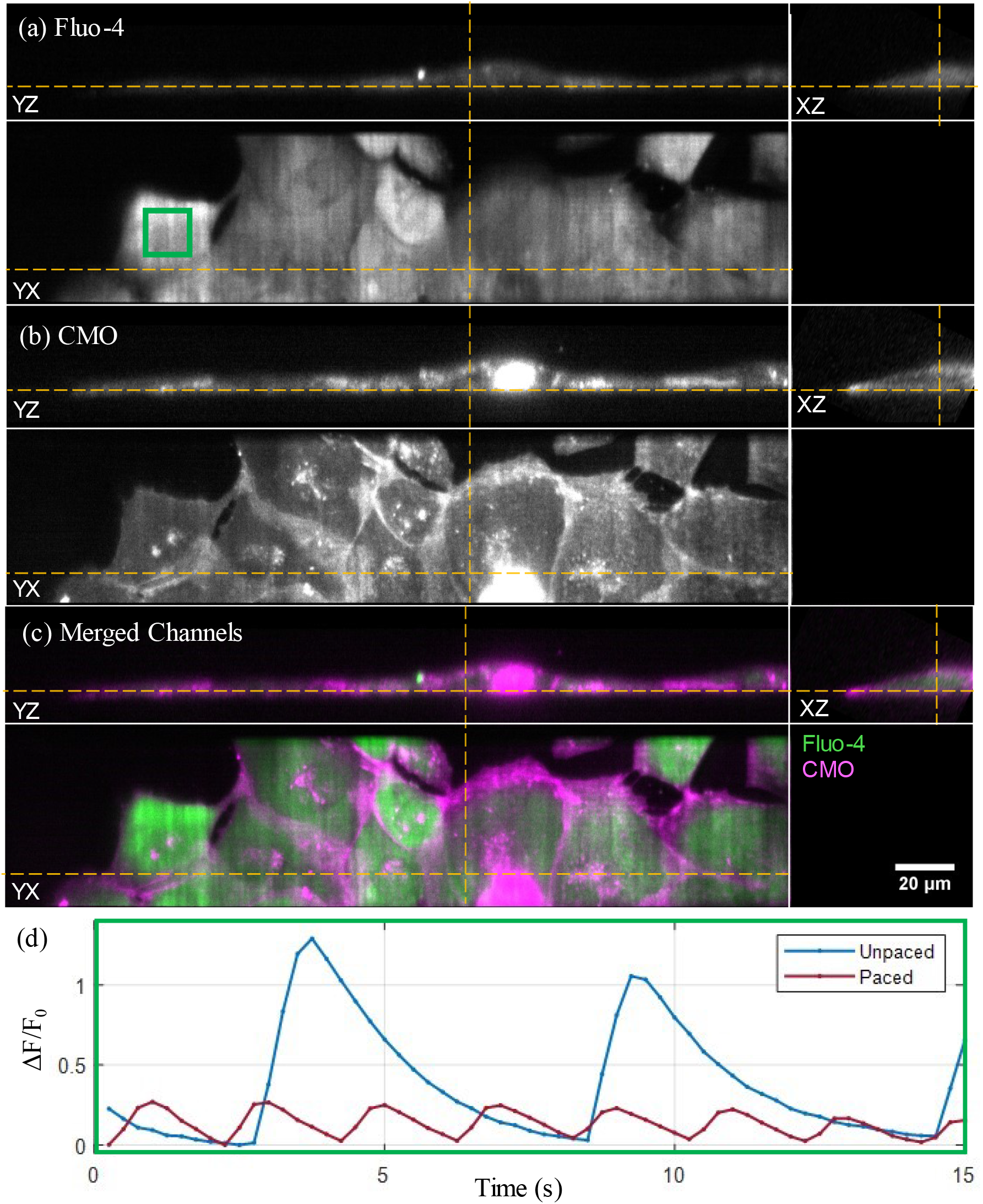
3D LSFM imaging of spontaneous and stimulated calcium transients in hiPSC-CM. Orthogonal cuts through the (a) Fluo-4, (b) CMO and (c) merged channels of hiPSC-CM culture imaged in 3D at 4 vps, 195 fps, displayed at the peak of the first spontaneous transient (t = 3.75 s) during an unpaced acquisition – the corresponding timelapse is shown in **Supplementary Video 1**. The yellow dashed lines indicate the intersecting planes of the displayed orthoslices. (d) Calcium transient ΔF/F_0_ time traces calculated over the three-dimensional 100 × 100 × 1 -voxel ROI (green rectangle in (a)) in consecutive unpaced and paced (at 0.5 Hz) acquisitions of the same FOV.

### Electromechanical coupling in hiPSC-CM and adult-CM co-culture

When observing the hiPSC-CM and adult-CM co-culture after 1 day, it was discovered that the added adult-CM exhibited spontaneous contraction without external electrical stimulation. These spontaneous dynamics were synchronized both with other adult-CM, and the beating of the underlying hiPSC-CM layer. This synchronization of transients and/or contraction between individual adult-CM and hiPSC-CM cells, here referred to as *coupling*, was investigated using widefield transillumination and fluorescence microscopy, and 3D LSFM.

Two co-cultures (A & B) were created on day 29 and day 34 respectively after start of hiPSC-CM differentiation. For each co-culture, a total of three samples were prepared (A_I-III_ & B_I-III_). For one of the three samples for each co-culture (A_I_ & B_I_), the stem cells were plated directly on the bottom of the well chamber for compatibility with widefield imaging. The co-culture development was observed on days 0, 1 and 2 after the start of the co-culture, with 3D volumetric LSFM imaging done on the plastic coverslip-plated samples A_II_, B_II_ and B_III_ on days 1&2.

### Widefield imaging of unpaced synchronized transients and contraction

Example transillumination images of samples co-cultured for 0-2 days are shown in **Figure 3**, with the corresponding timelapse presented in **Supplementary Video 2**. In the day 0 sample (a), the adult-CM can be distinguished from the underlying layer of hiPSC-CM by their rectangular shape and stronger scattering. After 24 hours of co-culture (b), the adult-CM are more rounded, and by day 2 of co-culture (c), most have a round shape. Unpaced, yet synchronized contractions between spatially separated adult-CM were observed as early as 4 hours after co-culture (a), with the cells exhibiting shortening along their long axis at the same time. Simultaneous beating of the underlying hiPSC-CM layer can also be observed, indicating that synchronization is occurring between the two cell types. Contracting cells can be identified from the brighter regions in the standard deviation maps for the widefield transillumination channel, shown in **Figure 3(d-f)**. For longer co-culture durations, as the adult-CM lose their elongated shape, distinguishing between their contraction and movement with the beating hiPSC-CM layer becomes more challenging, and hence unstimulated synchronized calcium transients are used as the primary evidence of coupling.

**Figure 3.**
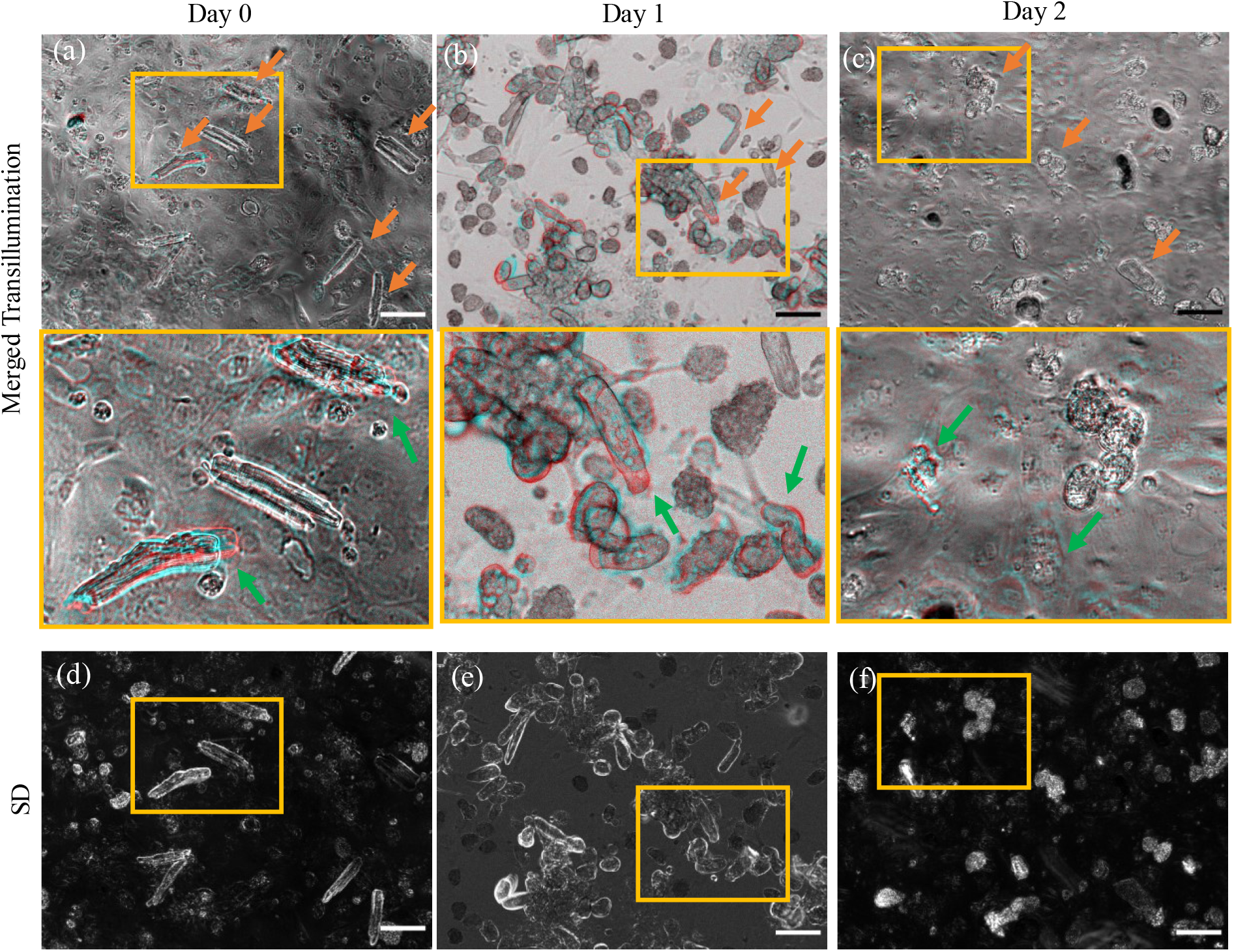
Synchronized contraction between spatially separated adult-CM in day 0, 1, 2 co-culture imaged under widefield transillumination. (a-c) Top images show two merged frames: at peak of contraction (cyan), and at baseline (red), taken t = 2.5 s apart for samples on (a) day 0, (b) day 1 and (c) day 2 of co-culture. Orange arrows indicate some of the adult cardiomyocytes on top of the layer of hiPSC-CM co-culture. (a-c) Bottom row shows zoomed in ROI indicated by the yellow rectangles in the top row, with green arrows pointing out some of the contraction movement between the merged frames (see **Supplementary Video 2** for 50 s timelapse). (d-f) Images for days 0, 1, and 2, calculated as the standard deviation across the whole acquisition, indicating movement during acquisition. The yellow rectangles indicate the zoomed in ROI shown in the bottom row of (a-c). Scalebar: 100 µm.

**Figure 4** shows examples from widefield epifluorescence recordings acquired on day 0, 1 and 2 of co-culture (left, middle and right columns respectively), with the corresponding timelapse presented in **Supplementary Video 3**. The fluorescence intensity distribution at baseline and at the peak of the first calcium transient is shown in the top and middle rows respectively, with the third row displaying the standard deviation in intensity across the acquisition duration. On day 0, some of the adult-CM (distinguishable from the underlying hiPSC-CM layer by their shape and difference in baseline intensity) undergo synchronized transients with the stem cells (examples are indicated by orange arrows). By day 1 and day 2 of co-culture, the majority of adult-CM have simultaneous transients with the hiPSC-CM. Examples of synchronized transients within individual adult-CM and areas of hiPSC-CM are shown through the ΔF/F_0_ time traces shown in the bottom row of **Figure 4** (panels k-m).

**Figure 4.**
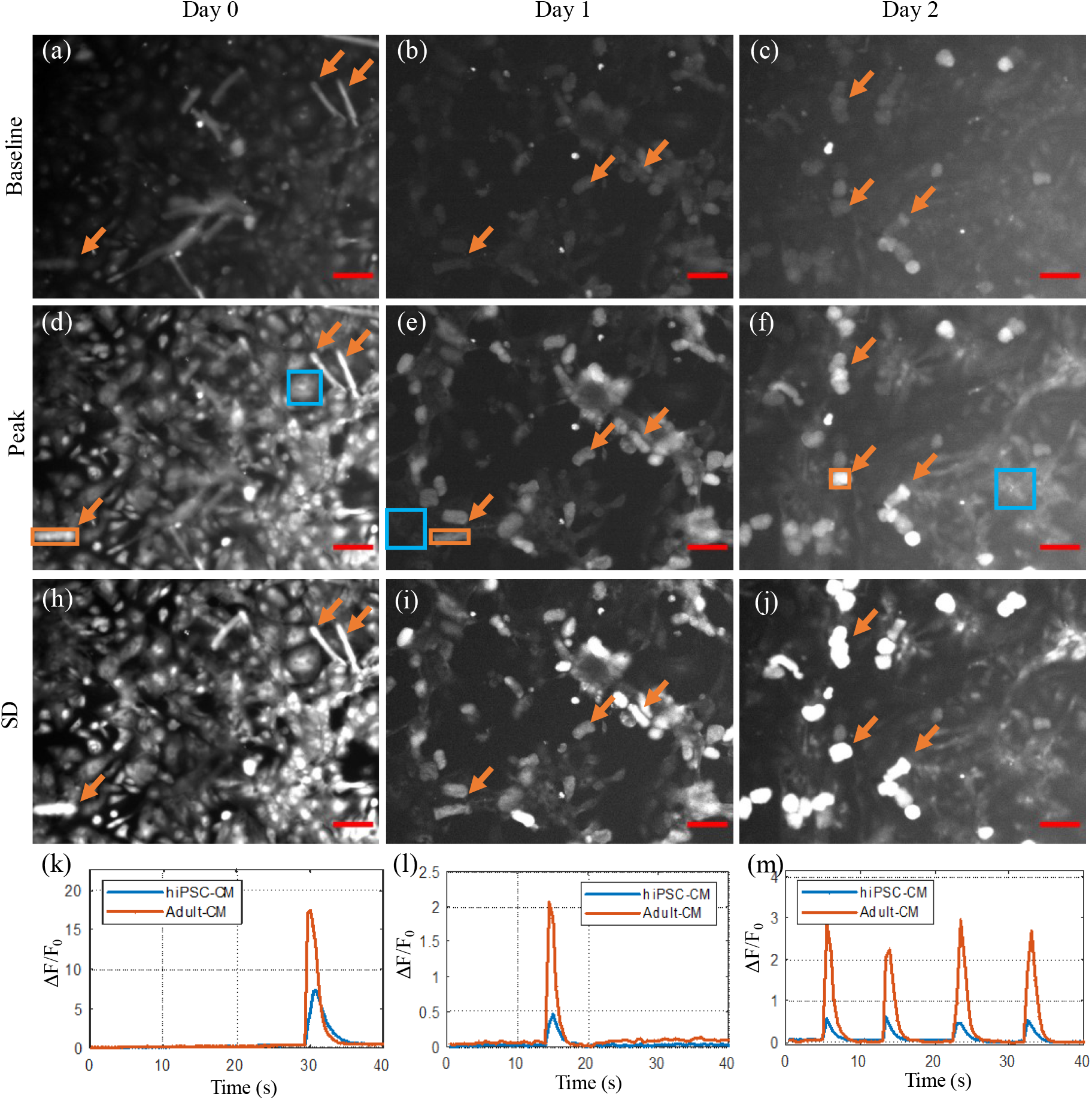
Widefield fluorescence imaging of calcium transients in adult-CM and hiPSC-CM co-culture on day 0 (left), day 1 (middle) and day 2 (right). Single frames at baseline (a-c) and peak of first transient (d-f), with frames from the same sample displayed on the same intensity scale. (h-j) Standard deviation calculated across the whole acquisition. Orange arrows indicate examples of coupled adult-CM. (k-m) Fluorescence intensity variation through hiPSC-CM (blue square ROI in middle row) and adult-CM (orange square ROI in middle row) indicating synchronized calcium transients. Scalebar: 100 µm. The corresponding timelapse is shown in **Supplementary Video 3**.

**Supplementary Table 3** summarizes the dynamics observed in day 0, 1 and 2 samples from two co-cultures with contraction and transient synchronicity observed in all samples except for one, which did not appear to have any visible dynamics – potentially due to poor sample health.

### 3D LSFM imaging of synchronized transients and contraction

Three of the samples from co-cultures A & B prepared on transferable plastic coverslips (A_II_, B_II_ and B_III_) were imaged in 3D on days 1 and 2 days of co-culture. For each of the 3 samples, ≥ 5 co-culture FOVs containing unique cardiomyocytes were imaged, with and without electrical pacing at 0.5 Hz. The optically sectioned 3D data allowed manual selection of ROI within each cell, hence enabling analysis of changes in fluorescence emission within that region, with minimal contribution from out-of-focus fluorescence.

**Figure 5** and **Supplementary Video 4a** demonstrate an example 3D time-lapse acquired at 8 Hz through a FOV within an unpaced day 1 co-culture of hiPSC-CM and adult-CM, with the corresponding depth-encoded MIP along the z-axis shown in Supplementary Video 4b. The adult-CM has retained an elongated shape, with some loss of rectangularity. By considering the variation of ΔF/F_0_ with time through ROIs selected for the hiPSC-CM and adult-CM (blue and orange respectively in panel (b)), it is possible to observe the transient synchronicity, with lower transient amplitude and slower decay in the hiPSC-CM.

**Figure 5.**
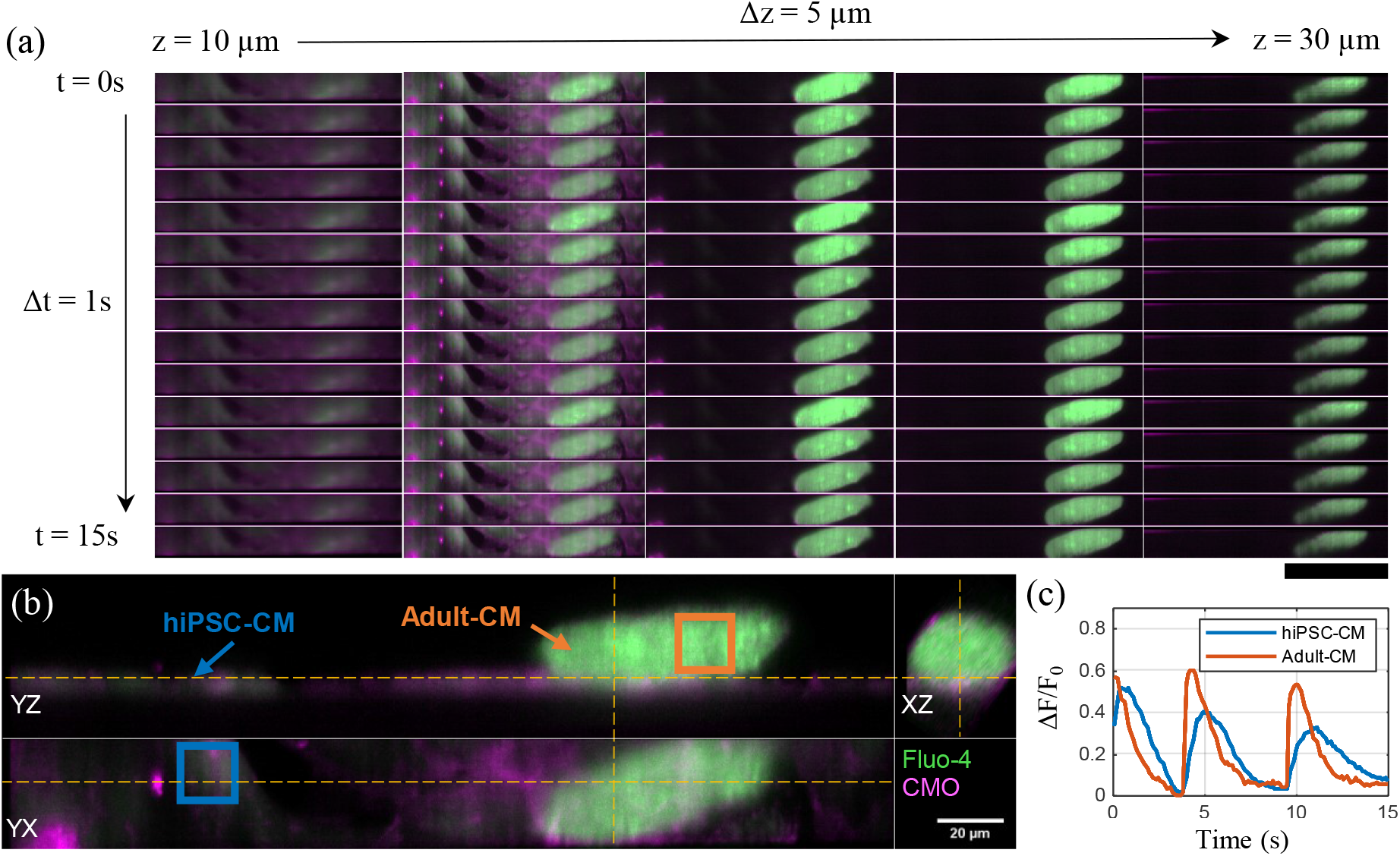
3D timelapse imaging of synchronized spontaneous transients in adult-CM and hiPSC-CM co-cultured for 24 hours. (a) Montage showing merged Fluo-4 (green) and CMO (magenta) 3D time-lapse for a single unpaced 15 s acquisition acquired *at* 8 vps, see **Supplementary Video 4a&b**. Each column represents XY-planes spaced 5 µm apart axially within the central third of the z-range (increasing upwards from coverslip) of the volume. Each row represents timepoints 1 s apart. Scalebar: 100 µm. (b) Orthogonal cuts through the merged channels displayed at the peak of the second transient (t =4.6 s). Yellow lines indicated the intersecting planes of the displayed orthoslices. (c) Calcium transient ΔF/F_0_ time traces calculated over the three-dimensional 100 × 100 × 1-pixel ROI within hiPSC-CM and adult-CM cells, indicated by the blue and orange rectangles respectively in (b).

The results over a total of 21 unique FOV, each containing a different adult-CM, from 3 different samples of two unique co-cultures (A & B) are summarized in **Supplementary Table 4**. For the day 1 co-culture, 6 out of 10 imaged adult-CM had coupled dynamics with hiPSC-CM. For the day 2 co-culture samples, the imaged FOVs did not contain any coupled events, however, there were fewer paced and unpaced calcium transients overall, with slower dynamics, and hence it is possible that the lack of transients from some of the imaged adult-CM from day 2 samples was due to selection of poor-quality cells. Additionally, by day 2 of co-culture, healthy adult-CM were more difficult to identify during sample pre-find in transillumination due to their loss of elongated shape and loss of regular t-tubule structure (compared to cardiomyocytes on day 0 of co-culture).

The limited FOV in the x’-direction (parallel to the propagation of the light sheet) made location of adult-CM within the co-culture challenging, with many elongated adult-CM from the day 1 sample not being aligned with the long axis of the rectangular FOV. To address this limitation, the FOV extent along the x’-direction was doubled for the experiments described in the following section. Co-culture duration was limited to days 0-1 to simplify identification of adult-CM and allow a more detailed investigation of the early-stage coupling of the two cell types. To evaluate whether it is the mechanical contraction of hiPSC-CM that drives the coupling of the two cell types, the following experiment investigated samples with and without mechanical decoupling with para-nitroblebbistatin.

#### Co-culture duration and immobilization influence on coupling

3D LSFM and widefield fluorescence microscopy were used to investigate coupling of adult-CM and hiPSC-CM for variable co-culture duration, and the presence and absence of the mechanical decoupler para-nitroblebbistatin (NBleb). Two co-cultures (C and D) of hiPSC-CM and adult-CM from two male rats were created on days 34 and 40 respectively past the start of hiPSC-CM differentiation. For each of the two adult-CM and hiPSC-CM co-cultures, 4 samples were prepared, and first two samples were labelled and imaged using 3D LSFM on the same day after ∼4 hr of co-culture, while the remaining two were imaged the following day, ∼24 hours after co-culture start. Within each pair, the second coverslip sample was prepared and imaged with mechanical decoupling using NBleb, while the first was prepared and imaged without. Additionally, the four samples prepared for co-culture D were imaged on a widefield fluorescence microscope immediately prior to LSFM imaging, to assess adult-CM contraction and presence of transients on day 0 & 1 of co-culture, with and without NBleb.

#### Widefield imaging of transients and contraction

Each of the four samples from co-culture D was imaged in two consecutive 40 s, 200 frame acquisitions, first in transillumination mode, then in fluorescence mode, focused on the adult-CM layer. The total number of adult-CM was counted from the image averaged over all frames from the transillumination acquisition (**Supplementary Figure 7a**). Contracting adult-CM were identified from the image of the standard deviation of the transillumination acquisition divided by the average (**Supplementary Figure 7b**) as those with bright outlines of the cell ends. Adult-CM with transients were identified from the image calculated as the standard deviation of all frames from the fluorescence channel acquisition, divided by the average (**Supplementary Figure 7c**). **Supplementary Figure 7d** demonstrates the merging of (a-c), indicating which cells had transients, contractions or both during the consecutive transillumination and fluorescence acquisitions.

The number of cells with contractions, transients for NBleb-treated and control (without NBleb) samples on day 0 and day 1 of co-culture are summarized in **Supplementary Table 5**. Samples with NBleb had significantly fewer cells contracting than those not treated with NBleb: 2% compared to 22% respectively on day 0 (p = 0.0008, Chi-squared test), and 6% compared to 39% respectively on day 1 (p = 0.0007, Chi-square test). Despite the mechanical decoupling, the samples with and without NBleb had no significant difference in the fraction of cells with calcium transients: 34% compared to 36% respectively on day 0 (p = 0.84 (n/s), Chi-square test) and 38% compared to 41% respectively on day 1 (p = 0.71 (n/s), Chi-squared test). This result indicates successful decoupling with NBleb, without any significant impact on transient prevalence.

The fraction of adult-CM with contraction in cells without NBleb was significantly higher for day 1 samples, compared to those imaged on day 0 (39% compared to 22% respectively, p = 0.0092, Chi-squared test). For NBleb samples, the increased number of contracting cells by day 1 was not statistically significant (p = 0.29, Chi-squared test), however, it is worth noting the overall small number of contracting cells in samples treated with NBleb. For calcium transients, the increase between day 0 and day 1 was not statistically significant for both control (p = 0.31, n/s, Chi-squared test) and NBleb-treated samples (p = 0.89, n/s, Chi-squared test).

#### 3D LSFM imaging of coupling events

Samples from both co-cultures C & D were imaged with 3D LSFM on day 0 and day 1 of co-culture, with and without NBleb, with twice the FOV along the x’-axis compared to the results presented above, at the cost of a decrease in volume acquisition rate from 8 Hz to 4 Hz due to an upper limit of a maximum frame rate of 199 fps for the readout of 1024 pixels in the vertical direction of the camera sensor. For each of the 8 samples, ≥ 5 FOVs containing unique cardiomyocytes were imaged over two consecutive acquisitions: first without electrical stimulation, then with electrical pacing at 0.5 Hz. Each time-lapse acquisition was manually visualized as an image stack and through MIPs along the detection axis. The optically sectioned 3D data allowed manual selection of ROI within each cell, hence allowing analysis of changes in fluorescence emission within that region, with minimal contribution from out-of-focus fluorescence.

**Figure 6** shows examples of synchronized dynamics in hiPSC-CM and adult-CM co-culture samples with and without mechanical decoupling. **Figure 6a&b** show orthogonal cuts through a volume from 15 s duration acquisitions of adult-CM and hiPSC-CM day 1 co-culture without NBleb (a) and with NBleb (b), after co-registration, axial rescaling, and rotation into lab coordinates. The corresponding orthogonal slice and depth encoded MIP timelapses are displayed in **Supplementary Videos 5a&b**, 6a&bThree-dimensional 100 × 100 × 1 -voxel ROI within the orthogonal cuts were used to sample the Fluo-4 intensity variation with time within hiPSC-CM and adult-CM. As can be seen from the ΔF/F_0_ variation with time for the two ROI in unpaced acquisitions in samples without and with NBleb (**Figure 6c&d** respectively), adult-CM and hiPSC-CM undergo synchronized spontaneous transients without external electrical stimulation at periods of T = 5.5 s and T = 5 s respectively. In **Figure 6c**, an increase in the baseline fluorescence of the adult-CM can be observed, which may be either a photoinduced response to the light-dose or an independent effect due to poor calcium control in the adult cardiomyocyte. There is plenty of scope to reduce the light dose while maintaining both temporal resolution and sufficient signal to noise ratio. For the ΔF/F_0_ time-traces shown in **Figure 6d**, the transient decay slope for adult-CM is visibly steeper than that for hiPSC-CM.

**Figure 6.**
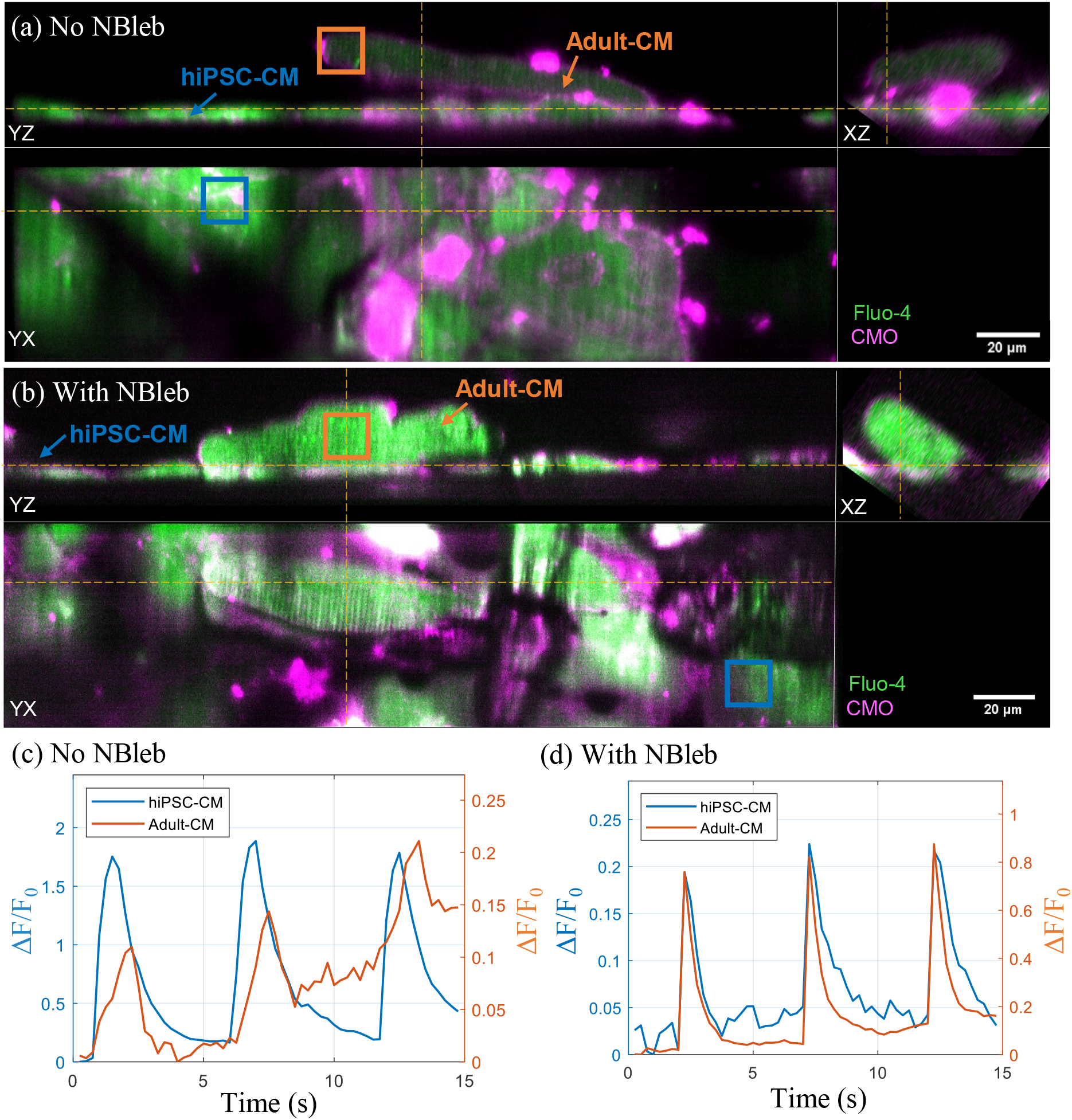
Synchronized spontaneous transients in co-culture in samples with and without NBleb. (a-b) Orthogonal cuts through the merged Fluo-4 (green) and CMO (magenta) channels of hiPSC-CM and adult-CM co-culture imaged in 3D at 4 vps, 195 fps during unpaced acquisitions for samples (a) without NBleb and (b) with NBleb displayed at the peak of the first spontaneous transient (t = 2 s and t = 2.5 s for (a) and (b) respectively). The corresponding time-lapses are shown in **Supplementary Video 5a&b, 6a&b**. Yellow lines indicated the intersecting planes of the displayed orthoslices. (c-d) Calcium transient ΔF/F_0_ time-traces calculated over the three-dimensional 100 × 100 × 1-voxel ROI within hiPSC-CM (blue) and adult-CM (orange) as indicated by the orange and blue rectangles respectively in (a) and (b).

An example of day 0 adult-CM coupling in co-culture is demonstrated in **Supplementary Figure 8** (and corresponding **Supplementary Videos 7a&b**), with two adult-CM within the same 3D LSFM FOV. While both adult-CM show activity in the form of calcium sparks, only the adult-CM on the left undergoes an unpaced transient, synchronously with the hiPSC-CM, while the right adult-CM only experiences a minor increase in baseline ΔF/F_0_ (**Supplementary Figure 8c)**. The 3D information in these acquisitions enables us to see that the right adult-CM comes very close to the hiPSC-CM layer, but despite this they are not coupled.

**Table 1** summarizes the samples prepared and imaged with 3D LSFM. Both unpaced and paced acquisitions for each FOV were manually screened for cell contraction either by hiPSC-CM or adult-CM, presence of transients in hiPSC-CM and adult-CM and coupled events between the two cell types. Adult-CM were classed as coupled when calcium dynamics occurred synchronously in both the adult-CM and any of the underlying hiPSC-CM cells. For paced acquisitions, adult-CM were not classed as coupled if synchronous transients were only observed at the pacing frequency, due to the ambiguity of whether the cells are coupled or responding to the applied stimulation.

**Table 1.**
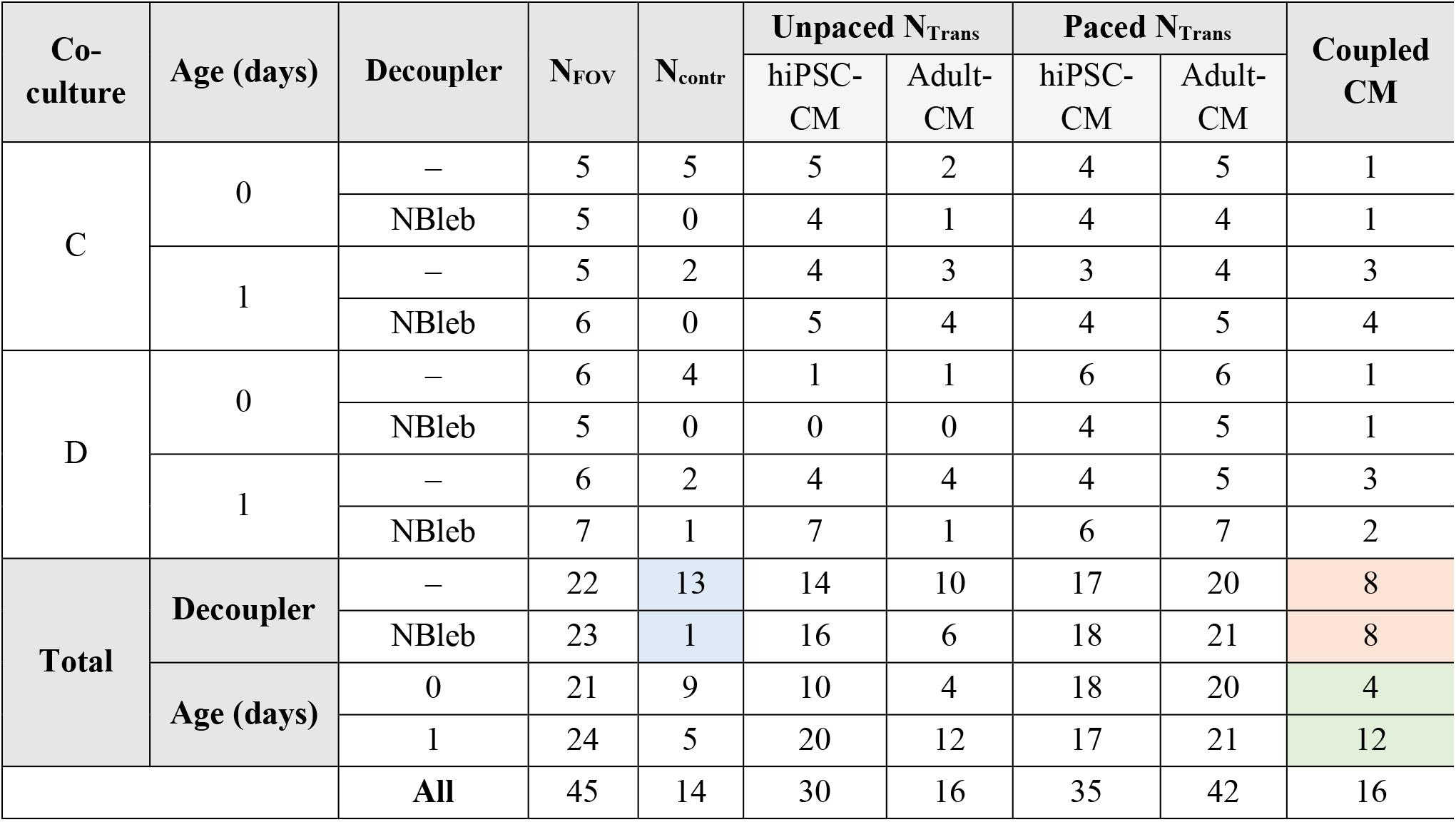
Summary of the 3D LSFM imaging of co-culture with and without NBleb. For each of the two co-cultures (referred to as C & D), two samples were imaged on the starting day of co-culture (day 0), while the other two were imaged the following day (day 1). Decoupling with para-nitroblebbistatin is indicated by “ Bleb”, with “–” indicating no decoupler used. The number of unique FOVs containing adult-CM imaged within each co-culture sample is indicated by N_FOV_, and N_contr_ indicates how many FOVs with those adult-CM contained cells contracting during the acquisition. The next six columns indicate how many of the N_CM_ contained hiPSC-CM and adult-CM underwent calcium transients in unpaced vs paced image acquisitions. Finally, the last column indicates how many out of the N_CM_ for each sample are considered coupled according to the criteria discussed above. While there were more adult-CM contracting in control samples, compared to those treated with NBleb (blue highlight) the number of coupled adult-CM was the same (red highlight). Overall, the number of coupled CM was higher by day 1 of co-culture, compared to day 0 (green highlight).

Across the two co-cultures, a total 45 FOVs with unique adult-CM co-cultured with hiPSC-CM were imaged, with each one imaged with and without electrical stimulation. As can be seen from **Table 1**, for samples treated with NBleb, only 1 out of 23 (4%) FOVs contained contracting cells, compared to 13 out of 22 FOVs (59%) from samples without NBleb (p = 0.0001, Chi-square test), indicating successful decoupling and in agreement with the results from widefield imaging. Overall, there was no significant difference in the number of adult-CM with coupled dynamics in samples with and without decoupler: 8 out of 22 FOVs (36%) from samples without NBleb, compared to 8 out of 23 FOVs in samples with NBleb (p = 0.91, n/s, Chi-square test), which indicates that the mechanical decoupling did not prevent synchronization of calcium dynamics in the two cell types.

Synchronized events were observed in samples co-cultured for only 4 hours, with 4 out of 21 CM (19%) imaged on the day the co-culture was started being classified as coupled. Additionally, in agreement with earlier observations on increased coupling between hiPSC-CM and adult-CM with longer co-culture time, coupling improved after 24 hours of co-culture, with 12 out of the 24 CM (50%) imaged in samples co-cultured for 24 hours (i.e. day 1) being classified as coupled, significantly higher than the proportion for day 0 (p = 0.0305, Chi-square test).

## Discussion

We have demonstrated a novel application of a custom light-sheet fluorescence microscope to volumetric imaging of calcium dynamics in live hiPSC-CM culture, and multi-layer co-culture with rat adult ventricular cardiomyocytes. Aberration-free remote refocusing of the detection plane synchronously to the scanning of the light sheet enabled acquisition of three-dimensional dual-channel optically sectioned timelapse data at up to 8 Hz with sub-cellular resolution over a ∼300 µm × 40 µm × 40 µm volume without any mechanical sample disturbance. As a result, it was possible to visualize and characterize the synchronized calcium dynamics and electromechanical coupling between hiPSC-CM and adult-CM, and quantify the prevalence of coupled events in samples co-cultured for different durations, with and without mechanical uncoupling using NBleb.

Using 2D and 3D LSFM imaging, the hiPSC-CM were observed to undergo both spontaneous unpaced beating and paced calcium transients and contraction at a range of pacing periods (*T* = 2 s, 4 s, 6 s). Complementing 3D LSFM imaging with larger FOV widefield fluorescence and transillumination microscopy of hiPSC-CM co-culture with adult ventricular cardiomyocytes isolated from n = 4 rats allowed the visualization of synchronized contraction and calcium transients in stem cell-derived and adult cardiomyocytes. The observation of synchronization as early as 4 hours after co-culture start was unexpected, since, for example, suspensions of individual hiPSC-CM condensed into engineered heart tissue take 10-15 days to couple with each other and create a beating construct [28]. A previous study considered the coupling of mouse iPSC-CM and mouse primary cardiomyocytes [17], however, coupling was only studied after 3 days, and the primary cells were the more plastic neonatal cardiomyocytes, rather than the adult-CM. In the present study, the coupling was progressive with time with significantly more adult cardiomyocytes acquiring the rhythm of the spontaneously beating hiPSC-CM after 24 hours of co-culture than on day 0.

This rapid establishment of coupling is difficult to reconcile with gap junction formation, given the predicted time for gene induction, protein expression and maturation of the connexin assembly [29]. We therefore examined other mechanisms for hiPSC-CM/adult cell synchronisation, such as mechanotransduction from the beating hiPSC-CM inducing calcium transients in the quiescent adult cardiomyocytes. However, while contraction in most cells was inhibited in the presence of a mechanical decoupler, the addition of NBleb did not prevent synchronization of calcium transients between adult cardiomyocytes and hiPSC-CM. Further experiments would be required to establish the exact coupling mechanism(s). The rate of gap junction formation between hiPSC-CM and adult-CM can be evaluated by imaging gap junction puncta in live cells [30], or by employing gap junction uncouplers such as carbenoxolone [31]. The possible role of ephaptic coupling could also be explored: this is where electrical excitation might be communicated between cardiomyocytes through e.g. an electric field effect, when gap junction transmission is not sufficient to maintain the action potential propagation [32].

A key limitation in this study was the sample size: for biologically distinct co-cultures, ventricular cardiomyocytes were sourced from a total of n = 4 rat hearts, with simultaneous access to both freshly isolated cells and hiPSC-CM at equivalent stages of differentiation presenting a logistical challenge due to their periodic availability. In future investigations, exploring the coupling interaction between species-matched iPSC-CM and adult-CM (human iPSC-CM/human adult-CM from failing heart explants, or rat iPSC-CM/isolated adult-CM) would be of interest.

A further experimental constraint was the co-culture duration, limited by the adult-CM experiencing loss of defined structure and elongated shape as they de-differentiated. Cardiomyocytes are known to undergo both morphological changes (including the loss of rod-like shape and t-tubule structure and reduction of cell size) and modifications in electromechanical behaviour (such as reduced systolic shortening and amplitude of calcium transients) in monoculture, and this is thought to be due to the loss of mechanical forces they are subjected to in the intact tissue. While these changes are a non-apoptotic and non-necrotic adaptation to the culture conditions [33] these phenotypical alterations limit the duration that the cultured cardiomyocytes can reliably be used as a model. Improved maintenance of the original morphology and function of cultured cardiomyocytes can be achieved with continuous electric field stimulation [34].

Another limitation was the 3D LSFM data acquisition and analysis throughput due to manual sample pre-find and acquisition, restricted FOV and minimally automated analysis. Low-NA widefield microscopy enables imaging of both calcium transients and contraction of over one hundred adult-CM in co-culture with hiPSC-CM. However, 2D imaging without optical sectioning prevented definite axial localization of the source of the fluorescence emission within the multi-layer co-culture. The optical sectioning of 3D LSFM reduced out-of-focus blur otherwise present in widefield imaging and allowed the origin of the fluorescence to be localized in 3D, with a single calcium indicator fluorophore simultaneously loaded into both cell types. Here, the combination of the two imaging methods allowed the coupling observed by widefield microscopy to be independently verified with 3D LSFM at subcellular resolution.

## Conclusion

Using a novel LSFM and low-NA widefield microscopy it was possible to visualize and characterize the synchronized calcium dynamics and electromechanical coupling between hiPSC-CM and adult-CM, quantifying the prevalence of coupled events in samples co-cultured for different durations, with and without mechanical uncoupling. The findings supported increased coupling with longer co-culture duration, with more adult-CM coupled to hiPSC-CM after 24 hours of co-culture (4 out of 21 (19%) vs 12 out of 24 (50%) on days 0 and 1 respectively, p = 0.0305). There was no significant difference between the number of adult-CM classed as coupled in samples treated with contraction uncoupler para-nitroblebbistatin compared to untreated samples, (8 out of 23 (35%) and 8 out of 22 (36%) and adult-CM respectively, p = 0.91), indicating that coupling is maintained in the absence of mechanotransduction.

In context of stem-cell regenerative therapy, the synchronized calcium dynamics and contraction in the heterogenous co-culture would be a desired outcome for optimum cardiac contraction. However, as shown by the co-culture model, the spontaneous beating of hiPSC-CM can drive calcium transients and contraction in the usually quiescent adult-CM once they are coupled. This behaviour could be potentially pro-arrhythmic, resulting in local activation and re-entry phenomena. Further research would be needed to evaluate how susceptible hiPSC-CM are to electrical stimulation at faster frequencies closer to the human heart rate of 0.6-2 Hz, and whether stimulated dynamics in hiPSC-CM dominate over or conflict with spontaneous beating.

While these results are more proof-of-concept rather than a comprehensive study, they demonstrate the power of the microscopical techniques described to answer important questions about stem cell coupling to the heart. The co-culture and imaging assay developed as part of this work can be used to investigate the influence of other environmental parameters and drugs on coupling efficiency. In the future, the coupling and interaction of hiPSC-CM and adult-CM can be studied using a model more closely representing stem cell grafting in vivo, with the iPSC-CM cells applied to living myocardial slice [35] to probe the coupling and synchronization of the graft cells with directly stimulated cardiac tissue [36]. The flexible remote-refocusing light-sheet fluorescence microscope may find future application in tackling the challenges of imaging dynamics in these thicker three-dimensional samples.

## Acknowledgements

The authors gratefully acknowledge funding from British Heart Foundation (BHF) (NH/16/1/32447). Liuba Dvinskikh acknowledges an Engineering and Physical Sciences Research Council (EPSRC) and BHF co-funded PhD studentship (EP/L015498/1) and Liliana Brito was the recipient of a PhD studentship funded by the BHF Centre of Research Excellence grant (RE/18/4/34215) The authors would like to thank Peter O’Gara for expert technical assistance in the isolation of ventricular cardiomyocytes, and Thusharika Kodagoda for differentiation of hiPSC-CM.

## Author contributions

L.D., S.H., and C.D. designed the experiments. H.S. developed the 3D remote-refocusing LSFM system and wrote the acquisition software. L.B. maintained and replated the hiPSC-CM culture. L.D. implemented the dual-channel detection system used on the LSFM, prepared the co-culture samples, performed the imaging experiments and data analysis. L.D. drafted the manuscript and prepared the figures. L.D., H.S., L.B., S.H., K.M., C.D. critically reviewed and contributed to the manuscript.

## Data availability

The raw data underlying this publication will be made freely available on request from the corresponding author*.

## References

[1] Bergmann, O. et al. Dynamics of cell generation and turnover in the human heart. Cell 161, 1566–1575, DOI: 10.1016/j.cell.2015.05.026 (2015).

[2] Breckwoldt, K., Weinberger, F. & Eschenhagen, T. Heart regeneration. Biochimica et Biophysica Acta (BBA) - Molecular Cell Research 1863, 1749–1759, DOI: 10.1016/j.bbamcr.2015.11.010, (2016).

[3] Kikuchi, K. & Poss, K. D. Cardiac Regenerative Capacity and Mechanisms. Annu. Rev. Cell Dev. Biol. 28, 719–741, DOI: 10.1146/annurev-cellbio-101011-155739 (2012).

[4] Eschenhagen, T. et al. Cardiomyocyte Regeneration. Circulation 136, 680–686, DOI: 10.1161/CIRCULATIONAHA.117.029343 (2017).

[5] Fisher, S. A. et al. Stem cell therapy for chronic ischaemic heart disease and congestive heart failure. Cochrane Database of Systematic Reviews, DOI: 10.1002/14651858.CD007888.pub3 (2014).

[6] Passier, R., van Laake, L. W. & Mummery, C. L. Stem-cell-based therapy and lessons from the heart. Nature 453, 322–329, DOI: 10.1038/nature07040 (2008).

[7] Chong, J. J. H. et al. Human embryonic-stem-cell-derived cardiomyocytes regenerate non-human primate hearts. Nature 510, 273–277, DOI: 10.1038/nature13233 (2014).

[8] Takahashi, K. & Yamanaka, S. Induction of Pluripotent Stem Cells from Mouse Embryonic and Adult Fibroblast Cultures by Defined Factors. Cell 126, 663–676, DOI: 10.1016/j.cell.2006.07.024 (2006).

[9] Morizane, A. et al. Direct comparison of autologous and allogeneic transplantation of iPSC-derived neural cells in the brain of a nonhuman primate. Stem cell reports 1, 283–292, DOI: 10.1016/j.stemcr.2013.08.007 (2013).

[10] Shiba, Y. et al. Allogeneic transplantation of iPS cell-derived cardiomyocytes regenerates primate hearts. Nature (London) 538, 388–391, DOI: 10.1038/nature19815 (2016).

[11] Kim, J. J. et al. Mechanism of automaticity in cardiomyocytes derived from human induced pluripotent stem cells. J. Mol. Cell. Cardiol. 81, 81–93, DOI: 10.1016/j.yjmcc.2015.01.013 (2015).

[12] Kane, C., Couch, L. & Terracciano, C. M. N. Excitation–contraction coupling of human induced pluripotent stem cell-derived cardiomyocytes. Front. Cell Dev. Biol. 3, DOI: 10.3389/fcell.2015.00059 (2015).

[13] Rao, C. et al. The effect of microgrooved culture substrates on calcium cycling of cardiac myocytes derived from human induced pluripotent stem cells. Biomaterials 34, 2399–2411, DOI: 10.1016/j.biomaterials.2012.11.055 (2013).

[14] Zhang, G. Q., Wei, H., Lu, J., Wong, P. & Shim, W. Identification and characterization of calcium sparks in cardiomyocytes derived from human induced pluripotent stem cells. PloS one 8, e55266, DOI: 10.1371/journal.pone.0055266 (2013).

[15] Lee, Y. et al. Calcium homeostasis in human induced pluripotent stem cell-derived cardiomyocytes. Stem Cell Reviews and Reports 7, 976–986, DOI: 10.1007/s12015-011-9273-3 (2011).

[16] Hwang, H. S. et al. Comparable calcium handling of human iPSC-derived cardiomyocytes generated by multiple laboratories. J. Mol. Cell. Cardiol. 85, 79–88, DOI: 10.1016/j.yjmcc.2015.05.003 (2015).

[17] Aratyn-Schaus, Y. et al. Coupling primary and stem cell–derived cardiomyocytes in an in vitro model of cardiac cell therapy. J. Cell Biol. 212, 389–397, DOI: 10.1083/jcb.201508026 (2016).

[18] Huisken, J., Swoger, J., Del Bene, F., Wittbrodt, J. & Stelzer, E. H. Optical Sectioning Deep Inside Live Embryos by Selective Plane Illumination Microscopy. Science 305, 1007–1009, DOI: 10.1126/science.1100035 (2004).

[19] Sparks, H. et al. Development a flexible light-sheet fluorescence microscope for high-speed 3D imaging of calcium dynamics and 3D imaging of cellular microstructure. J Biophotonics 13, e201960239, DOI: 10.1002/jbio.201960239 (2020).

[20] Botcherby, E. J., Juškaitis, R., Booth, M. J. & Wilson, T. An optical technique for remote focusing in microscopy. Opt. Commun. 281, 880–887, DOI: 10.1016/j.optcom.2007.10.007 (2008).

[21] Botcherby, E. J., Juskaitis, R., Booth, M. J. & Wilson, T. Aberration-free optical refocusing in high numerical aperture microscopy. Opt. Lett., OL 32, 2007–2009, DOI: 10.1364/OL.32.002007 (2007).

[22] Jabbour, R. J. et al. In vivo grafting of large engineered heart tissue patches for cardiac repair. JCI insight 6, DOI: 10.1172/jci.insight.144068 (2021).

[23] Sato, M., O’Gara, P., Harding, S. E. & Fuller, S. J. Enhancement of adenoviral gene transfer to adult rat cardiomyocytes in vivo by immobilization and ultrasound treatment of the heart. Gene Ther. 12, 936–941, DOI: 10.1038/sj.gt.3302476 (2005).

[24] Sikkel, M. B. et al. High speed sCMOS-based oblique plane microscopy applied to the study of calcium dynamics in cardiac myocytes. J Biophotonics 9, 311–323, DOI: 10.1002/jbio.201500193. (2016).

[25] Képiró, M. et al. para-itroblebbistatin, the non-cytotoxic and photostable myosin II inhibitor. Angewandte Chemie International Edition 53, 8211–8215, DOI: 10.1038/sj.gt.3302476 (2014).

[26] Katrukha, E. (2021) Z-stack Depth Color Code ImageJ/Fiji Plugin https://github.com/ekatrukha/ZstackDepthColorCode

[27] Germanguz, I., et al., Molecular characterization and functional properties of cardiomyocytes derived from human inducible pluripotent stem cells. J Cell Mol Med. 15 (1), 38–51. DOI: 10.1111/j.1582-4934.2009.00996.x (2011)

[28] Breckwoldt, K. et al. Differentiation of cardiomyocytes and generation of human engineered heart tissue. Nature protocols 12, 1177–1197, DOI: 10.1038/nprot.2017.033 (2017).

[29] van Veen, T.A.B., van Rijen, H.V.M., Opthof, T., Cardiac gap junction channels: modulation of expression and channel properties. Cardiovascular Research 51(2), 217–229, DOI: 10.1016/S0008-6363(01)00324-8 (2001).

[30] Katoch, P. et al. The Carboxyl Tail of Connexin32 Regulates Gap Junction Assembly in Human Prostate and Pancreatic Cancer Cells. The Journal of biological chemistry 290, 4647–4662, DOI: 10.1074/jbc.M114.586057 (2015).

[31] de Groot, J. R. et al. Conduction slowing by the gap junctional uncoupler carbenoxolone. Cardiovascular Research 60, 288–297, DOI: 10.1016/j.cardiores.2003.07.004 (2003).

[32] Sperelakis, N. An electric field mechanism for transmission of excitation between myocardial cells. Circulation Research 91 985–987 DOI: 10.1161/01.res.0000045656.34731.6d (2002).

[33] Banyasz, T. et al. Transformation of adult rat cardiac myocytes in primary culture. Exp. Physiol. 93, 370–382, DOI: 10.1113/expphysiol.2007.040659 (2008).

[34] Berger, H. et al. Continual electric field stimulation preserves contractile function of adult ventricular myocytes in primary culture. American Journal of Physiology-Heart and Circulatory Physiology 266, H341–H349, DOI: 10.1152/ajpheart.1994.266.1.H341 (1994),.

[35] Watson, S. A., Terracciano, C. M. & Perbellini, F. Myocardial slices: an intermediate complexity platform for translational cardiovascular research. Cardiovascular Drugs and Therapy 33, 239–244, DOI: 10.1007/s10557-019-06853-5 (2019).

[36] Chen C., Bai X., Ding Y., Lee I., Electrical stimulation as a novel tool for regulating cell behavior in tissue engineering. Biomater. Res. 23(25) DOI: 10.1186/s40824-019-0176-8 (2019)

