## Supplementary Information for "Remote-refocusing light-sheet fluorescence microscopy enables 3D imaging of electromechanical coupling of hiPSC-derived and adult cardiomyocytes in co-culture"

### Methods – sample preparation

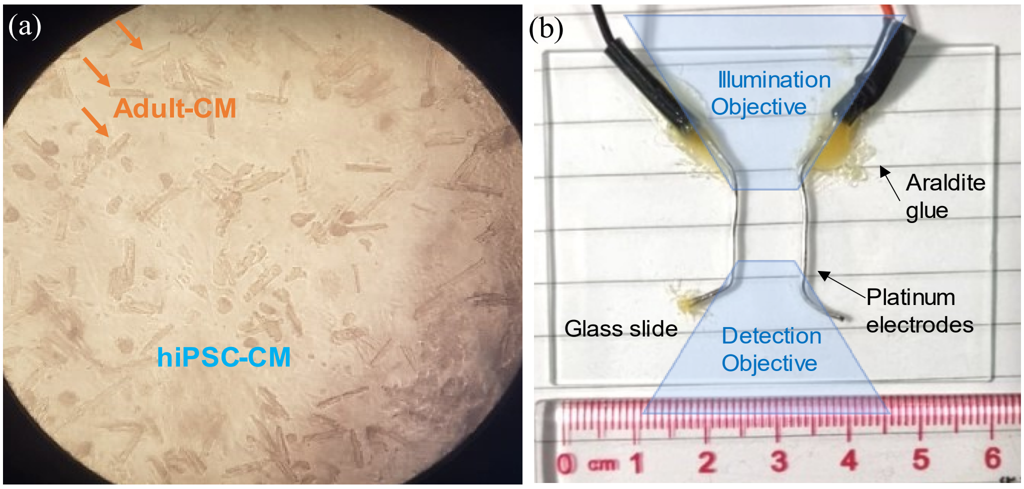


**Supplementary Figure 1. hiPSC-CM and adult-CM co-culture and pacing chamber.** (a) Representative distribution of adult-CM on top of a weakly scattering layer of hiPSC-CM, as seen ~4 hours after co-culture start. Image recorded through the eyepiece of an inverted transillumination widefield microscope through the glass bottom of the well and the plastic coverslip using a 4× air objective. (b) Top view of the chamber used for imaging and electrical stimulation of hiPSC-CM and their co-culture with adult-CM. Approximate objective positioning is illustrated in blue.

### Methods – experiment overview

| Exp. | Sample | Mode | FOV (x-y-z,µm) | FPS | VPS | Duration (s) | Pacing Period (s) | Co-culture duration (days) | Decoupl. |
|---|---|---|---|---|---|---|---|---|---|
| 1 | hiPSC-CM | 2D | 170×38 | 175 | N/A | 57 | –/2/4/6 | N/A | – |
| 2 | hiPSC-CM | 3D | 302×76×48 | 195 | 4 | 15 | –/2 | N/A | – |
| | | | 302×38×48 | 390 | 8 | 15 | | | |
| 3 | Co-cultures A & B | 3D | 302×38×48 | 390 | 8 | 15 | –/2 | 0, 1, 2 | –/NBleb |
| 4 | Co-cultures C & D | 3D | 302×76×48 | 195 | 4 | 15 | –/2 | 0, 1 | –/NBleb |

**Supplementary Table 1. Overview of the main time-lapse experiments with hiPSC-CM cells and their co-culture with adult LV CM,** and the key parameters and acquisition settings: the sample type, acquisition mode (2D or 3D), FOV dimensions (per spectral channel), frame and volume acquisition rate in frames per second (FPS) and volumes per second (VPS) respectively, total duration of each acquisition, presence and period of the electrical pacing, co-culture duration and presence of motion decoupler (NBleb). Lack of pacing or decoupler is denoted by "–".

### Methods – dual-channel imaging

#### Supplementary note 1 - correction of defocus in the Fluo-4 spectral channel

Small amounts of defocus can be generated by introducing two lenses with near-equal but opposite magnification, separated by a small distance. Using the thin lens approximation, the total optical power $K_{12} = {}^{1}/_{f_{12}}$ of two lenses with powers $K_1$ and $K_2$ is given by the Gullstrand equation [1]:

$$K_{12} = K_1 + K_2 - d_{12} K_1 K_2 \qquad (1)$$

where $d_{12}$ is the distance between the principal planes of the two lenses. As $d_{12}$ increases, the effective power of the two lenses decreases. The combined power of the lens pair and the 200mm tube lens (T4 in [2]) in front of the detector is given by

$$K_{tot} = K_{12} + K_T - d_T K_{12} K_T \qquad (2)$$

Where $K_T = {}^{1}/_{200\,mm}$ is the optical power of the tube lens, and the distance from approximate combined principal plane of the lens pair to the principal plane of tube lens (T3 in [2]) was estimated to be around $d_T$ =125 mm.

The defocus correction was achieved with two cemented doublet lenses $L_+$ and $L_-$ with focal lengths of $f_1$ = 160 mm and $f_2$ = -150 mm respectively (**Supplementary Fig. 2**), introduced within the Fluo-4 beam path of the dichroic assembly, after the emission filter and before positioning mirror (F1 and M12 respectively in **Fig. 1** in [2])

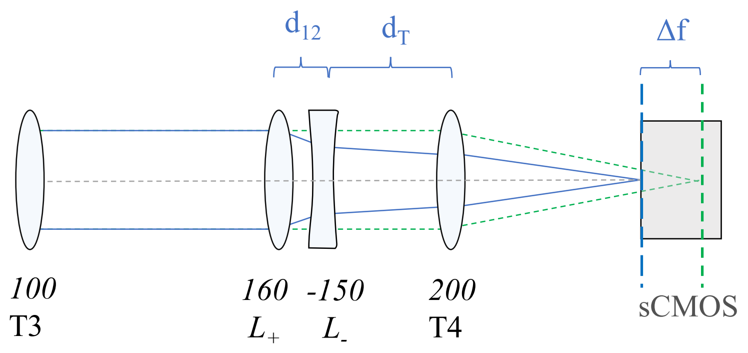


**Supplementary Figure 2.** Optical relay from tube lens T3 to the detector in the Fluo-4 emission path. The blue rays show how lenses $L_+$ and $L_-$ with principal planes separated by $d_{12}$~15mm introduce a Δf~10mm negative defocus compared to the original ray path (green). All focal lengths are given in mm.

Optical modelling in Zemax OpticStudio was used to determine the lens principal plane location, and optimal order and orientation (see Section 2.2.1 in Ref [3]). For the selected lens orientation with the positive lens positioned first, the theoretical principal plane separation of $d_{12} = 2.79$ mm, the expected nominal positive focus shift was $\Delta f = 12.8$ mm, which decreased with increasing lens separation. The required defocus correction of $\Delta f = -10$ mm is achieved at around $d_{12} = 15$ mm, which corresponds to a spacing of ~12 mm between the adjacent surfaces of the two lenses. After inserting the lens pair into the Fluo-4 beam path, Fluo-4 channel defocus was corrected by adjusting the lens pair spacing to ~1 cm. As a result, the focus for the Fluo-4 channel now coincided with that for the CMO channel, however, this introduced a magnification factor into the Fluo-4 channel. The resultant change in lateral magnification for the Fluo-4 channel was measured to be equal to M = 1.08 and was corrected for during the channel co-registration in post-processing.

### Methods – 3D LSFM imaging of hiPSC-CM and co-culture

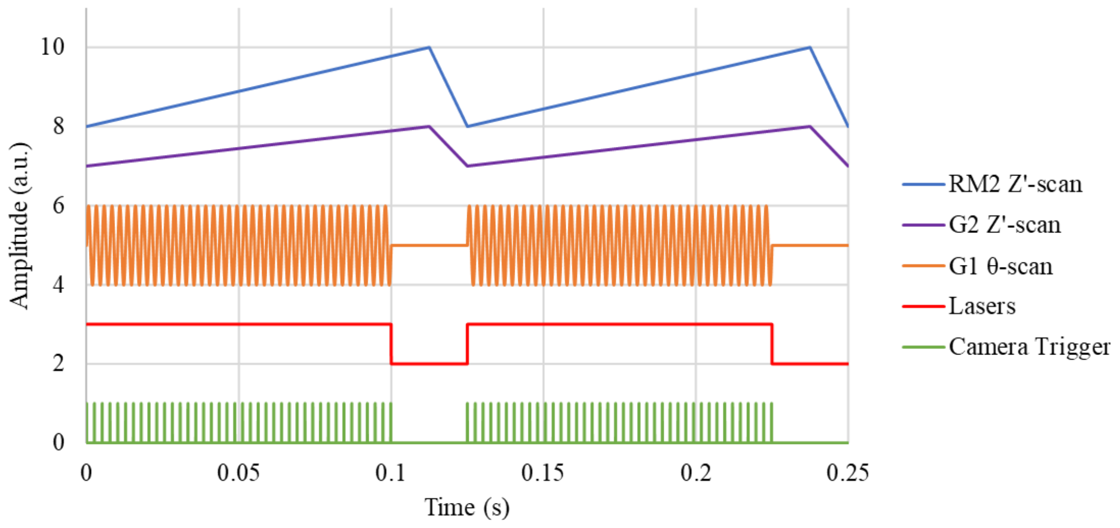


**Supplementary Figure 3. Hardware timing diagram for volumetric acquisition with remote refocus.** The diagram shows the output signal generated in the LabVIEW acquisition software for two consecutive volumes, each containing 40 planes, acquired at 8 Hz (~400 fps). Through the DAQ card, the camera and lasers were triggered through TTL signals, and analog output signals were used to control the z-scan galvo G1, dither galvo G2 and remote refocusing mirror RM2 positions. The amplitudes for each signal have been offset vertically for clearer visualization.

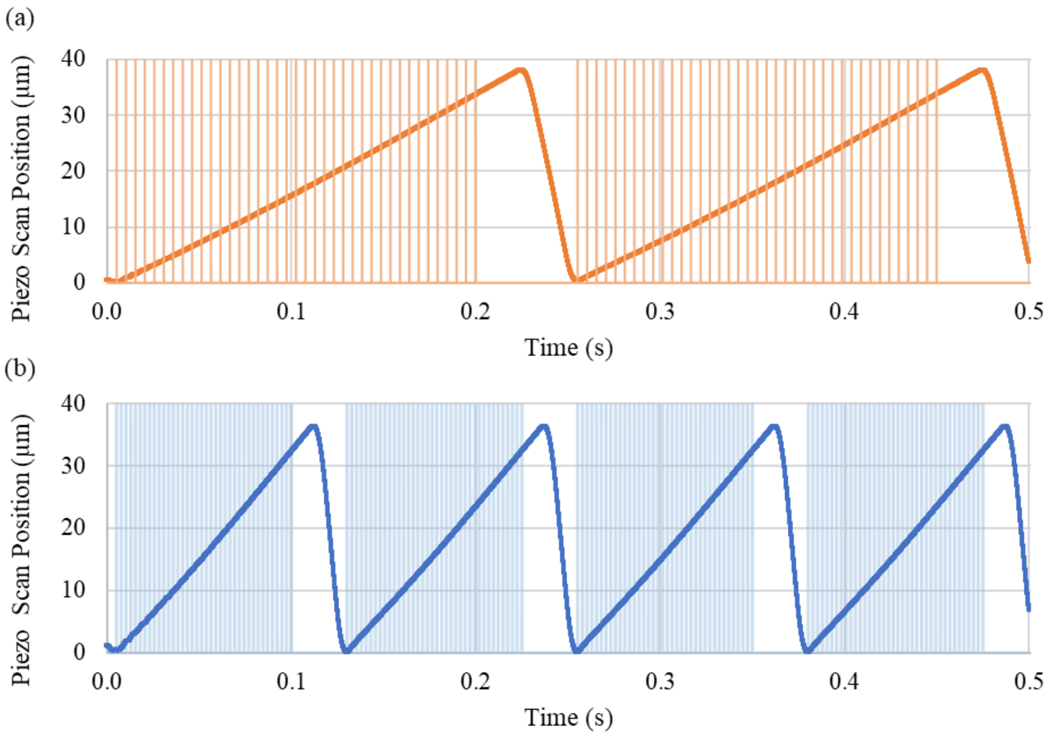


**Supplementary Figure 4. Exemplar recorded scan position of the piezo actuator used for remote-refocusing of the detection plane** for (a) 4Hz acquisition rate and (b) 8 Hz acquisition rate, with the camera frames within the acquisition window indicated in light orange (a) and light blue (b). The acquisition window is limited to the linear region of the piezo scan range, resulting in a ~33 µm scan range in remote-refocus space, which, when accounting for the double pass of the folded remote refocus and 1.33× magnification to sample space, is equal to a ~48 µm scan range in sample space.

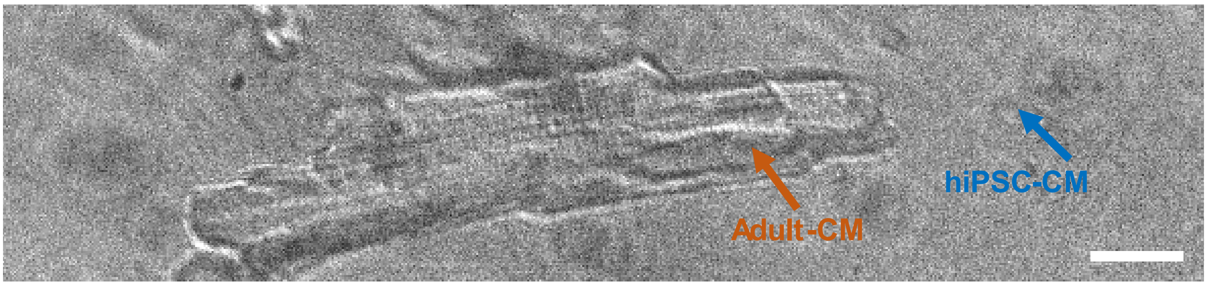


**Supplementary Figure 5.** Adult-CM on top of weakly-scattering hiPSC-CM, as seen in transillumination on the light sheet fluorescence microscope. Scalebar: 20 µm.

### Results – hiPSC-CM imaging with 2D LSFM

#### Supplementary note 2 - 2D LSFM imaging of spontaneous and stimulated transients in hiPSC-CM

2D timelapse imaging of FOV's from n = 3 different hiPSC-CM culture samples was carried out on separate occasions at 30 days past start of differentiation (**Supplementary Table 2**). Out of 19 unique imaged FOVs within the cultures with calcium transients, 11 acquisitions were electrically stimulated at varying periods: $T = 2$ s, 4 s and 6 s, and mechanical contraction of the cells was identified by visual screening of each dataset. Out of 19 unique imaged FOVs within the cultures with calcium transients, 11 acquisitions were electrically stimulated at varying periods: $T = 2$ s, 4 s and 6 s, and mechanical contraction of the cells was identified by visual screening of each dataset.

| hiSPC-CM culture | No of FOVs with transients | No of paced FOV | Pacing period (s) | No of FOV with contraction |
|---|---|---|---|---|
| I | 6 | 0 | – | 6 |
| II | 4 | 3 | –/2 | 2 |
| III | 9 | 8 | –/2/4/6 | 6 |
| **Combined** | 19 | 11 | –/2/4/6 | 14 |

**Supplementary Table 2. Summary of 2D timelapse acquisitions of unpaced and paced hiPSC-CM culture**, imaged on three different occasions. All acquisitions consisted of 10,000 frames at 175 fps (57 s duration), over a 1152 × 512 px FOV, imaged with and without electrical pacing.

**Supplementary Figure 6** illustrates various calcium dynamics present in a selection of unpaced and paced 2D timelapse acquisitions from the three hiPSC-CM cultures. Calcium transient $\Delta F/F_0$ vs time traces were analyzed by defining square ROIs within the Fluo-4 channel (green). The CMO channel (magenta) allowed visualization of the cell membrane and boundaries between adjacent cells. Cell layers ranged from a single cell to localized centres with a thickness of >10 cells, with most acquisitions limited to imaging areas with up to ~5 cells layers.

In the absence of external electric field stimulation, spontaneous transients were observed with and without contraction, for periods ranging between 4-14 s. In **Supplementary Figure 6a**, unpaced hiPSC-CM have spontaneous transients at a period of $T \sim 12.5$ s. Electrically stimulated calcium transients occurred both with and without contraction (data not shown). In **Supplementary Figure 6b**, the cells are electrically stimulated at 0.5 Hz and produce calcium transients at the stimulation frequency. The signal amplitude does not plateau at a baseline and is lower than in the slower spontaneous dynamics in (a). For 2 of 5 acquisitions paced at $T = 2$ s, paired transients, separated by a longer delay between consecutive pairs, were observed, with an example demonstrated in **Supplementary Figure 6c.** Slower, more consistent stimulated transients were observed at a period of $T = 4$ s and 6 s (**Supplementary Figure 6d&e** respectively). Although we have not demonstrated here, the high frame rate in the 2D LSFM imaging mode would allow the delay between electrical stimulation and the transient onset to be mapped spatially with ms timing accuracy.

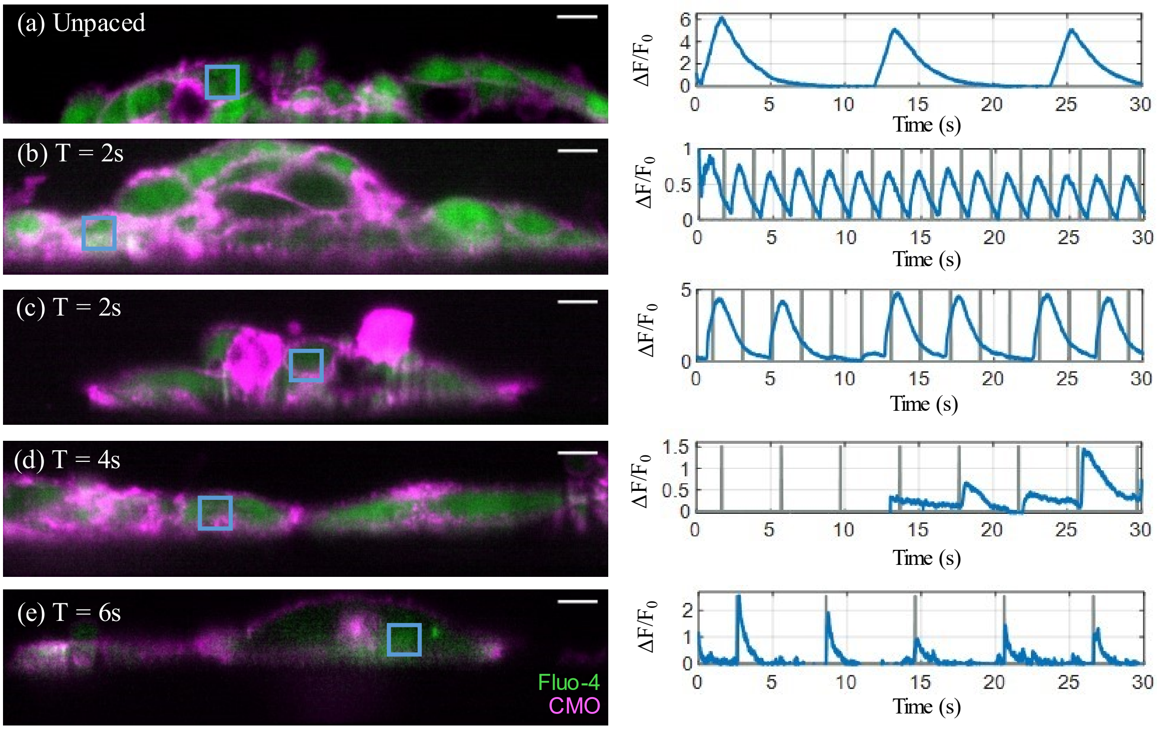


**Supplementary Figure 6. 2D imaging of spontaneous and stimulated calcium transients in hiPSC-CM.** Left column: merged Fluo-4 (green) and CMO (magenta) channels of a single frame at the peak of the first transient from a 10,000 frame (57 s) 2D time-lapse dataset imaged at 175 fps. Right column: corresponding Fluo4 $\Delta F/F_0$ time-traces (blue) taken through the 50 × 50 2D ROI indicated by the blue square outlines in the left column. The panels show cells unpaced (a) and paced at a period of (b) T = 2 s, (c) T = 2 s, (d) T = 4 s, and (e) T = 6 s, with a pulse duration of 2 ms. Panels (a), (b) and (c-e) show cells from the I, II and III hiPSC-CM culture samples respectively. The pacing timing is shown in grey. The dynamics were synchronized within most cells in the FOV. The first part of acquisition (d) lacks signal due to an unintentionally closed laser shutter. Scalebar: 10 µm.

### Results – co-culture, synchronized transients and contraction

| Co-culture | Age (days) | Sample | Widefield Imaging Observations | | | |
|---|---|---|---|---|---|---|
| | | | Transillum. – sync. contraction | | Fluoresc. – sync. transients | |
| | | | hiPSC-CM | Adult-CM | hiPSC-CM | Adult-CM |
| A | 1 | $A_{I}$ | + | + | + | + |
| | | $A_{II}$ | + | + | N/A | |
| | | $A_{III}$ | + | + | – | – |
| | 2 | $A_{II}$ | + | + | + | + |
| B | 0 | $B_{I}$ | + | +/– | + | +/– |
| | 1 | $B_{II}$ | + | + | N/A | |
| | | $B_{III}$ | + | + | +/– | +/– |
| | 2 | $B_{II}$ | +/– | +/– | + | +/– |

**Supplementary Table 3. Summary of dynamics identified from widefield transillumination and fluorescence imaging of the hiPSC-CM and adult-CM co-cultures A & B after 0, 1 and 2 days of co-culture**. Presence of synchronized contraction and transients of 5+ cells is denoted as "+", while the presence in <5 less is denoted as "+/–", and samples without identified synchronized transients or contraction are labelled as "–". Samples $A_{II}$ and $B_{II}$ were not labelled or imaged in fluorescence mode on day 1 and are labelled as N/A (grey).

| Co-culture | Age (days) | Sample | $N_{FOV}$ | Unpaced Transients | | | | Stimulated Transients | | | | Coupled CM |
|---|---|---|---|---|---|---|---|---|---|---|---|---|
| | | | | $N_{UnP}$ | hiPSC-CM | Adult-CM | Coupled Events | $N_P$ | hiPSC-CM | Adult-CM | Coupled Events | |
| A | 2 | $A_{II}$ | 6 | 3 | 2 | 0 | 0 | 5 | 5 | 3 | 0 | 0 |
| B | 1 | $B_{III}$ | 10 | 10 | 4 | 1 | 1 | 10 | 9 | 9 | 6 | 6 |
| | 2 | $B_{II}$ | 5 | 5 | 0 | 0 | 0 | 5 | 1 | 1 | 0 | 0 |
| **Total** | Age (days) | 1 | 10 | 10 | 4 | 1 | 1 | 10 | 9 | 9 | 6 | 6 |
| | | 2 | 11 | 8 | 3 | 0 | 0 | 11 | 6 | 1 | 0 | 0 |

**Supplementary Table 3. Summary of the 3D LSFM imaging of day 1 and day 2 co-cultures of hiPSC-CM and adult-CM A and B.** The number of unique FOVs containing adult-CM imaged within each co-culture sample is indicated by $N_{FOV}$. $N_{UnP}$ and $N_P$ indicate the number of unpaced and paced acquisitions respectively. The number of acquisitions containing transients in unpaced and paced acquisitions in both hiPSC-CM and adult-CM were counted. The last column indicates how many out of the $N_{FOV}$ for each co-culture sample contained adult-CM that were considered coupled according to the criteria discussed in the main text.

### Results – widefield imaging of co-culture with and without NBleb

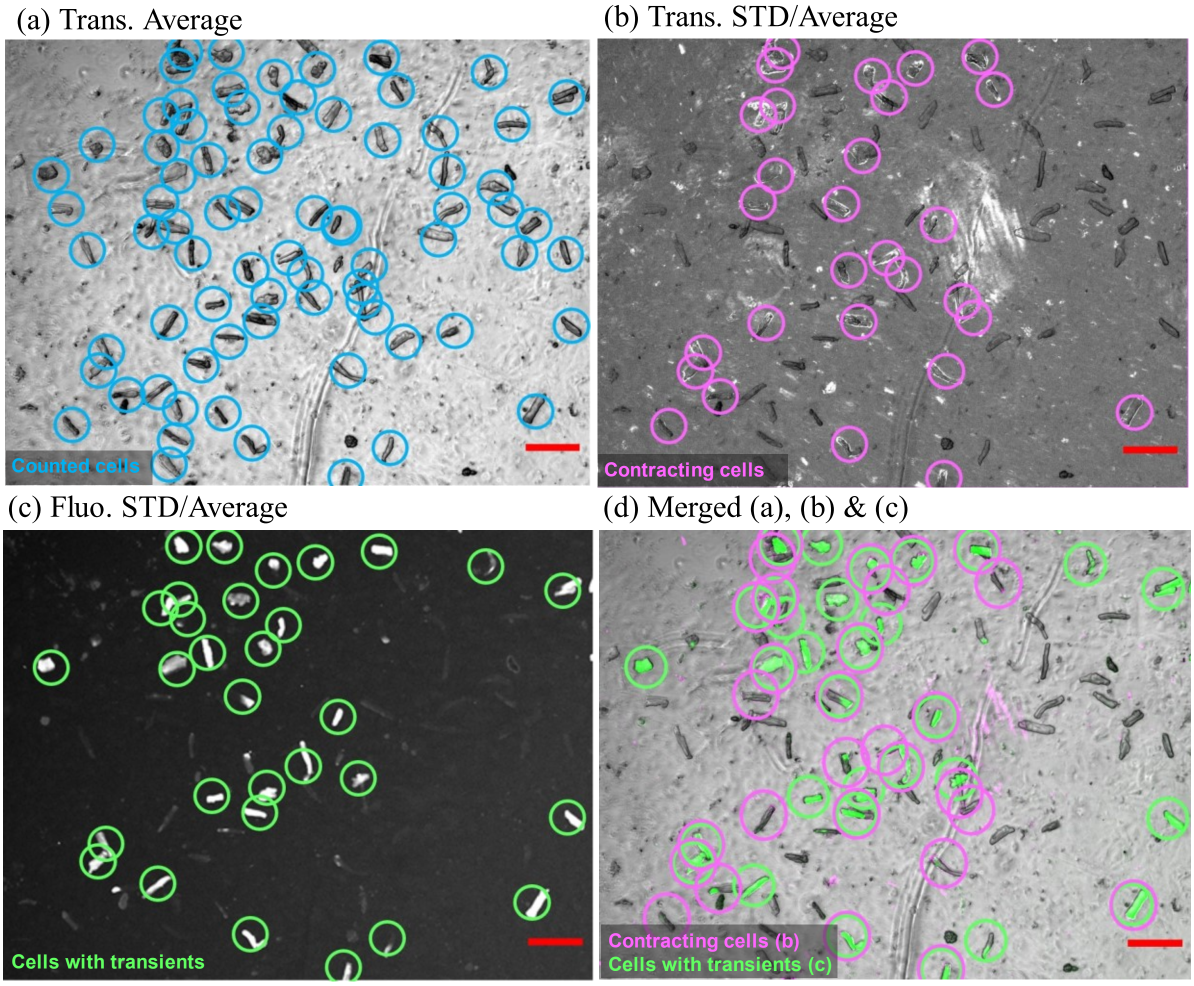


**Supplementary Figure 7. Counting adult-CM with transients and contraction from fluorescence and transillumination widefield images of co-culture.** Average of 200 frames (40 s duration acquisition at 5 fps) using (a) widefield transillumination and (b) and fluorescence (Fluo-4 channel) microscopy of the same FOV for day 1 no NBleb sample for co-culture D. Images (c) and (d) show the standard deviation divided by the average over the whole acquisition for the transillumination and fluorescence channels respectively. The merged images (a), (b) and (c) are shown in (d), in overlayed grey, magenta and green colours respectively. All counted adult-CM are encircled in blue in (a), those contracting in magenta (b and d) and those with transients in green (c and d). Scalebar: 200 μm.

| Co-culture | Age (days) | Decoupler | $N_{CM}$ | Transillumination | | Fluorescence | |
|---|---|---|---|---|---|---|---|
| | | | | $N_{contr}$ | % | $N_{Tr}$ | % |
| D | 0 | – | 137 | 30 | 22 | 47 | 34 |
| | | NBleb | 53 | 1 | 2 | 19 | 36 |
| | 1 | – | 75 | 29 | 39 | 31 | 41 |
| | | NBleb | 32 | 2 | 6 | 12 | 38 |

**Supplementary Table 4. Results from widefield transillumination and fluorescence imaging of day 0 and day 1 co-culture with and without NBleb.** Summary of observations from widefield transillumination and fluorescence imaging of the hiPSC-CM and adult-CM co-culture D after 4 and 24 hours of co-culture (day 0 and day 1) in samples prepared with and without decoupling ("–" and "NBleb" respectively). $N_{CM}$ indicates the counted number of adult-CM within the FOV, while $N_{contr}$ and $N_{Tr}$ indicate how many of these cells contract and have calcium transients respectively, with the percentage indicated in the adjacent column.

### Results – 3D LSFM Co-culture imaging

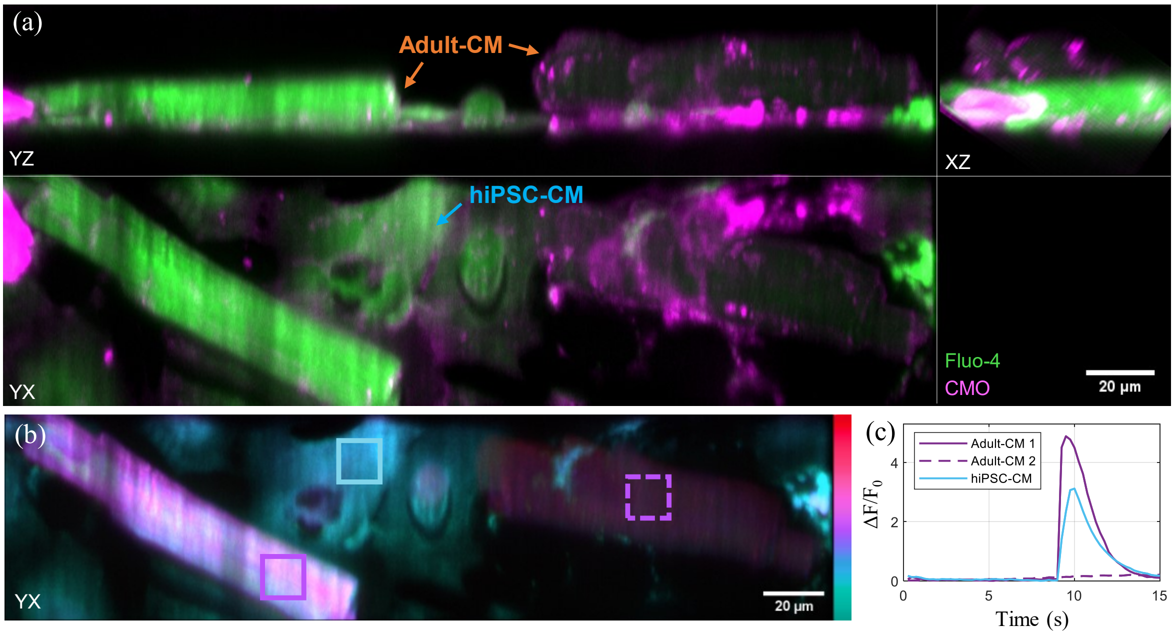


**Supplementary Figure 8. Synchronized spontaneous transient in one of the two adult-CM with the hiPSC-CM in day 0 co-culture sample without NBleb.** (a) Orthogonal maximum intensity projections (MIPs) through the merged Fluo-4 (green) and CMO (magenta) channels of hiPSC-CM and adult-CM co-culture imaged in 3D at 4 vps, 195 fps during unpaced acquisitions for samples without NBleb displayed at the peak of the spontaneous transient (t = 10 s). (b) Depth-encoded maximum intensity projection through the Fluo-4 channel along the z-axis over a 30 µm distance from the coverslip, displayed at the same timepoint as (a). Colorbar: 0 µm (cyan) – 30 µm (red). The outline colour of the 100 × 100 × 1-voxel ROIs indicates the depth of the plane of the ROI. The corresponding time-lapses are shown in **Supplementary Video 7a&b** [4]. (c) Calcium transient $\Delta F/F_0$ time-traces calculated over the ROIs within hiPSC-CM (blue), the left adult-CM (purple), and right adult-CM (purple, dashed) in (b).

### Video captions

The supplementary videos have been made available on Zenodo [4].

**Video 1: 3D LSFM timelapse of hiPSC-CM undergoing spontaneous calcium transients**. Orthogonal cuts through the merged Fluo-4 (green) and CMO (magenta) channels of hiPSC-CM culture imaged at 4 vps, 195 fps during an unpaced acquisition. Playback at 2× real-time speed.

**Video 2: Widefield transillumination timelapse of hiPSC-CM and adult-CM** day 0 (left), day 1 (middle) and day 2 (right) of co-culture, with synchronized spontaneous contraction of spatially separated adult-CM. Playback at 4× real-time speed. Scalebar: 100 µm.

**Video 3: Widefield fluorescence timelapse of hiPSC-CM and adult-CM** on day 0 (left), day 1 (middle) and day 2 (right) of co-culture, with synchronized spontaneous calcium transients. Corresponding $\Delta F/F_0$ time-traces through ROI of select adult-CM (orange) and hiPSC-CM (blue) are shown in **Figure 4k-m** in the main text. Playback at 4× real-time speed. Scalebar: 100 µm.

**Video 4a: 3D LSFM timelapse of hiPSC-CM and adult-CM day 1 co-culture undergoing synchronized spontaneous transients**. Orthogonal cuts through the merged Fluo-4 (green) and CMO (magenta) channels of hiPSC-CM culture, imaged at 8 vps, 390 fps during an unpaced acquisition. Corresponding $\Delta F/F_0$ time-traces through select ROI in adult-CM (orange) and hiPSC-CM (blue) are shown in **Figure 5c** in the main text. Depth-encoded MIP timelapse along the z-axis is shown in **Video 4b**. Playback at 2× real-time speed.

**Video 4b: Depth-encoded MIPs of the 3D LSFM timelapse of hiPSC-CM and adult-CM day 1 co-culture undergoing synchronized spontaneous transients**. Depth-encoded maximum intensity projection through the Fluo-4 channel imaged at 8 vps along the z-axis. The colorbar indicates distance up from the coverslip over a 30 µm range (0 µm (cyan) – 30 µm (red)). The corresponding orthogonal cut timelapse and $\Delta F/F_0$ time-traces through select ROI in adult-CM and hiPSC-CM are shown in **Video 4a** and **Figure 5** in the main text. Playback at 2× real-time speed.

**Video 5a: 3D LSFM timelapse of hiPSC-CM and adult-CM day 1 co-culture undergoing synchronized spontaneous transients** in a sample without NBleb. Orthogonal cuts through the merged Fluo-4 (green) and CMO (magenta) channels, imaged at 4 vps, 195 fps during an unpaced acquisition. Corresponding $\Delta F/F_0$ time-traces through select ROI in adult-CM (orange) and hiPSC-CM (blue) are shown in **Figure 6c** in the main text. Depth-encoded MIP timelapse along the z-axis is shown in **Video 5b**. Playback at 2× real-time speed.

**Video 5b: Depth-encoded MIPs of the 3D LSFM timelapse of hiPSC-CM and adult-CM day 1 co-culture without NBleb undergoing synchronized spontaneous transients**. Depth-encoded maximum intensity projection through the Fluo-4 channel imaged at 4 vps along the z-axis. The colorbar indicates distance up from the coverslip over a 30 µm range (0 µm (cyan) – 30 µm (red)). The corresponding orthogonal cut timelapse and $\Delta F/F_0$ time-traces through select ROI in adult-CM and hiPSC-CM are shown in **Video 5a** and **Figure 6a,c** in the main text. Playback at 2× real-time speed.

**Video 6a: 3D LSFM timelapse of hiPSC-CM and adult-CM co-culture undergoing synchronized spontaneous transients** in a sample treated with NBleb. Orthogonal cuts through the merged Fluo-4 (green) and CMO (magenta) channels, imaged at 4 vps, 195 fps during an unpaced acquisition. Corresponding $\Delta F/F_0$ time-traces through select ROI in adult-CM (orange) and hiPSC-CM (blue) are shown in **Figure 6d** in the main text. Depth-encoded MIP timelapse along the z-axis is shown in **Video 6b**. Playback at 2× real-time speed.

**Video 6b: Depth-encoded MIPs of the 3D LSFM timelapse of hiPSC-CM and adult-CM day 1 co-culture with NBleb undergoing synchronized spontaneous transients**. Depth-encoded maximum intensity projection through the Fluo-4 channel imaged at 4 vps along the z-axis. The colorbar indicates distance up from the coverslip over a 30 µm range (0 µm (cyan) – 30 µm (red)). The corresponding orthogonal cut timelapse and $\Delta F/F_0$ time-traces through select ROI in adult-CM and hiPSC-CM are shown in **Video 6a** and **Figure 6b,d** in the main text. Playback at 2× real-time speed.

**Video 7a: 3D LSFM timelapse of hiPSC-CM and adult-CM day 0 co-culture undergoing synchronized spontaneous transients** in a sample without NBleb. Orthogonal MIPs through the merged Fluo-4 (green) and CMO (magenta) channels, imaged at 4 vps, 195 fps during an unpaced acquisition. Corresponding $\Delta F/F_0$ time-traces through selected ROI in adult-CM and hiPSC-CM are shown in **Supplementary Figure 8.** Depth-encoded MIP timelapse along the z-axis is shown in **Video 7b**. Playback at 2× real-time speed.

**Video 7b: Depth-encoded MIPs of the 3D LSFM timelapse of hiPSC-CM and adult-CM day 0 co-culture without NBleb.** Depth-encoded maximum intensity projection through the Fluo-4 channel imaged at 4 vps along the z-axis. The colorbar indicates distance up from the coverslip over a 30 µm range (0 µm (cyan) – 30 µm (red)). The corresponding orthogonal cut timelapse and $\Delta F/F_0$ time-traces through select ROI in adult-CM and hiPSC-CM are shown in **Video 7a** and **Supplementary Figure 8** in the main text. Playback at 2× real-time speed.

### References

[1] Hecht, E. *Optics.* 4th edition. Addison-Wesley. (2002)

[2] Sparks, H. *et al*. Development a flexible light-sheet fluorescence microscope for high-speed 3D imaging of calcium dynamics and 3D imaging of cellular microstructure. *J Biophotonics* **13**, e201960239, DOI: 10.1002/jbio.201960239 (2020).

[3] Dvinskikh, L. Remote refocusing light-sheet fluorescence microscopy for high-speed 2D and 3D imaging of calcium dynamics in cardiomyocytes, Imperial College London, DOI: 10.25560/99343 (2022).

[4] Dvinskikh, L. *et al.* Supplementary videos for the "Remote-refocusing light-sheet fluorescence microscopy enables 3D imaging of electromechanical coupling of hiPSC-derived and adult cardiomyocytes in co-culture" manuscript (Version 1). Zenodo. DOI: 10.5281/zenodo.7580163. (2023)
